# Nucleic acid sensing by STING induces an interferon-like antiviral response in a marine invertebrate

**DOI:** 10.1101/2022.12.18.520954

**Authors:** Haoyang Li, Sheng Wang, Qinyao Li, Shaoping Weng, Jianguo He, Chaozheng Li

## Abstract

The cytosolic detection of pathogen derived nucleic acids has evolved as an essential strategy for host innate immune defense in mammals. The stimulator of interferon genes (STING) functions as a crucial signaling adaptor, linking the cytosolic detection of DNA by cyclic GMP-AMP (cGAMP) synthase (cGAS) to the downstream Type I interferon (IFN) signaling axis. However, this process remains elusive in invertebrates. Herein, we demonstrated that a STING ortholog from a marine invertebrate (shrimp) *Litopenaeus vannamei* can directly sense DNA to activate an interferon-like antiviral response. Unlike STING homologs exclusively functioning as a sensor for cyclic dinucleotides (CDNs) in other eukaryotic organisms, shrimp STING can bind to double-stranded DNA (dsDNA) in addition to CDNs, including 2′3′-cGAMP. *In vivo*, shrimp STING can directly sense DNA nucleic acids from an infected virus, accelerate IRF dimerization, nuclear translocation and induce the expression of an interferon functional analog protein (Vago4), and finally establish an antiviral state. Surprisingly, the shrimp cGAS-like homolog is not involved in dsDNA-intrigued and STING-dependent IRF–Vago axis activation. Taken together, our results uncovered a novel dsDNA–STING–IKKε–IRF–Vago antiviral axis in an arthropod, and provided some novel insights into the functional origin of a DNA-sensing pathway in evolution.

## INTRODUCTION

Animals are constantly facing challenges from pathogenic microbes. To counter infection, eukaryotic cells express pattern-recognition receptors (PRRs) to detect invading microbes through pathogen-associated molecular patterns (PAMPs) and activate innate immune responses [1]. As all microbial pathogens contain RNA or DNA or both for their replication, the recognition of microbial nucleic acids by various PRRs has evolved as a central defensive strategy of the innate immune system [2]. Three categories of innate immune sensors endosomal localized transmembrane Toll-like receptors (TLRs), cytosolic microbial RNA sensors, and cytosolic DNA sensors, are responsible for recognizing the nucleic acids of intracellular pathogens [3]. TLRs mainly detect various microbial DNA and RNA in the endosomal lumen [4]. Retinoic acid–inducible gene I (RIG-I)-like receptors, including RIG-I, MDA5, and LGP2, mainly sense pathogen-derived RNA in the cytosol [5]. Cyclic GMP-AMP (cGAMP) synthase (cGAS), a general cytosolic DNA sensor, mainly detects pathogen-derived DNA in the cytosol [6]. After binding to microbial nucleic acids, these innate immune sensors can initiate downstream signaling cascades that result in the production of type I interferon (IFN) or inflammatory cytokines or both [7].

Over the recent decade, innate immune responses elicited by cytosolic DNA derived from a large variety of DNA-containing pathogens have been intensively studied in mammals. Originally, a stimulator of interferon genes (STING, also known as TMEM173, MITA, ERIS and MPYS) is discovered to be indispensable for innate immune signaling in response to cytosolic DNA, as manifested by the loss of STING abrogates the ability of intracellular B-form DNA, as well as members of the herpesvirus family, to induce IFN-β [8–11]. Studies show that STING can be triggered by cyclic dinucleotides (CDNs) [12–14], cytosolic DNA [15] and DNA sensors including DEAD-box polypeptide 41 (DDX41), interferon-inducible protein 16 and meiotic recombination 11 homolog A [16–18]. To date, STING is a signaling mediator downstream of cGAS. cGAS are receptors with a zinc-ribbon domain for DNA binding, and biosynthetic enzymes with a nucleotidyltransferase domain, which catalyzes the synthesis of 2′3′-cyclic GMP-AMP (2′3′-cGAMP) from ATP and GTP [6, 19, 20]. cGAMP serves as a second messenger that binds to the endoplasmic-reticulum (ER)-resident protein STING [12]. cGAMP binding induces the oligomerization of STING at the ER-Golgi intermediate compartment and Golgi in the prenuclear region where STING recruits TBK1 via its C-terminal tail (CTT), which then results in TBK1 clustering and trans-autophosphorylation. Activated TBK1 phosphorylates STING C-terminal at the pLxIS motif (p, hydrophilic residue; x, any residue), allowing it to recruit IRF3, which in turn is phosphorylated by TBK1 [21, 22]. Phosphorylated IRF3 forms a dimer and translocates to the nucleus, and finally induces the transcription of IFN-ꞵ. In addition, NF-κB signaling output, the expression of inflammatory cytokines such as TNFα, IL-1ꞵ, and IL-6, is the observed downstream of STING activation [23]. Although the cGAS–STING pathway for cytosolic DNA sensing has been widely recognized in antiviral innate immunity in mammals, the function and regulation of the parallel pathway in invertebrates are still largely unknown.

In response to DNA species, vertebrate STING activation relies on 2′3′-cGAMP produced by cGAS [24]. Except *Crassostrea gigas* cGAS [25], no functional cGAS homolog with the ability to sense DNA in invertebrates has been reported [26]. Whether DNA can trigger a STING-dependent antiviral immune response in invertebrates remains unrevealed. Notably, the interferon antiviral system has long been considered a vertebrate specific immune pathway, our previous studies suggested that a parallel system exists in shrimp as well [27]. The antiviral system in shrimp shows remarkable similarities to the vertebrate interferon system, which has an IRF-like transcription factor, a secreted interferon-like cytokine (Vago) that activates a JAK/STAT signaling pathway and establishes an antiviral state [28, 29]. In the present study, we uncovered that shrimp *Litopenaeus vannamei* (Arthropoda: Crustacea: Decapoda) STING can bind to exogenous dsDNA directly, including virus-derived DNA, to activate an innate antiviral immunity through the IRF–Vago antiviral axis. Given that shrimp cGAS homolog might not participate in DNA-intrigued and STING-dependent signaling activation, we concluded that the direct binding to DNA by shrimp STING to activate innate immunity might bypass the need of a functional cGAS for DNA sensing, which can be an evolutionary innovation in the innate immune system in a marine invertebrate.

## RESULTS

### Identification and sequence analysis of LvSTING

Using human STING sequence as the query to BLAST against *L. vannamei* transcriptome data, we identified a sequence (GenBank No. KY490589) encoding a putative STING ortholog (named as LvSTING), which comprised 316 amino acids with an estimated molecular mass of 35.3 kDa. We then cloned the LvSTING genome, which contained nine exons and >23,590 bp of genomic DNA (Figure 1–figure supplement 1 A). The first two exons encoded 5’-untranslated sequence, exons 3–5 encoded four transmembrane domains (TMs) in the amino-terminal portion of LvSTING, and the last four exons combined to form the LBD and CTT domain (Figure 1–figure supplement 1A). In the analysis of sequence homology, STING homologs sequences from various species were retrieved from GenBank and used for phylogenetic analysis. The phylogenetic tree fell into two clades (Figure 1–figure supplement 1B). All STING homologs from vertebrates including *Homo sapiens* (HsSTING) formed a separate branch, whereas LvSTING and STING homologs from invertebrates including *Drosophila melanogaster* (DmSTING) were clustered together. The phylogenetic tree topology showed that LvSTING formed an isolated branch from phylum arthropod STING homologs, suggesting that LvSTING plays a crucial role during STING homologs evolution.

**Figure 1.**
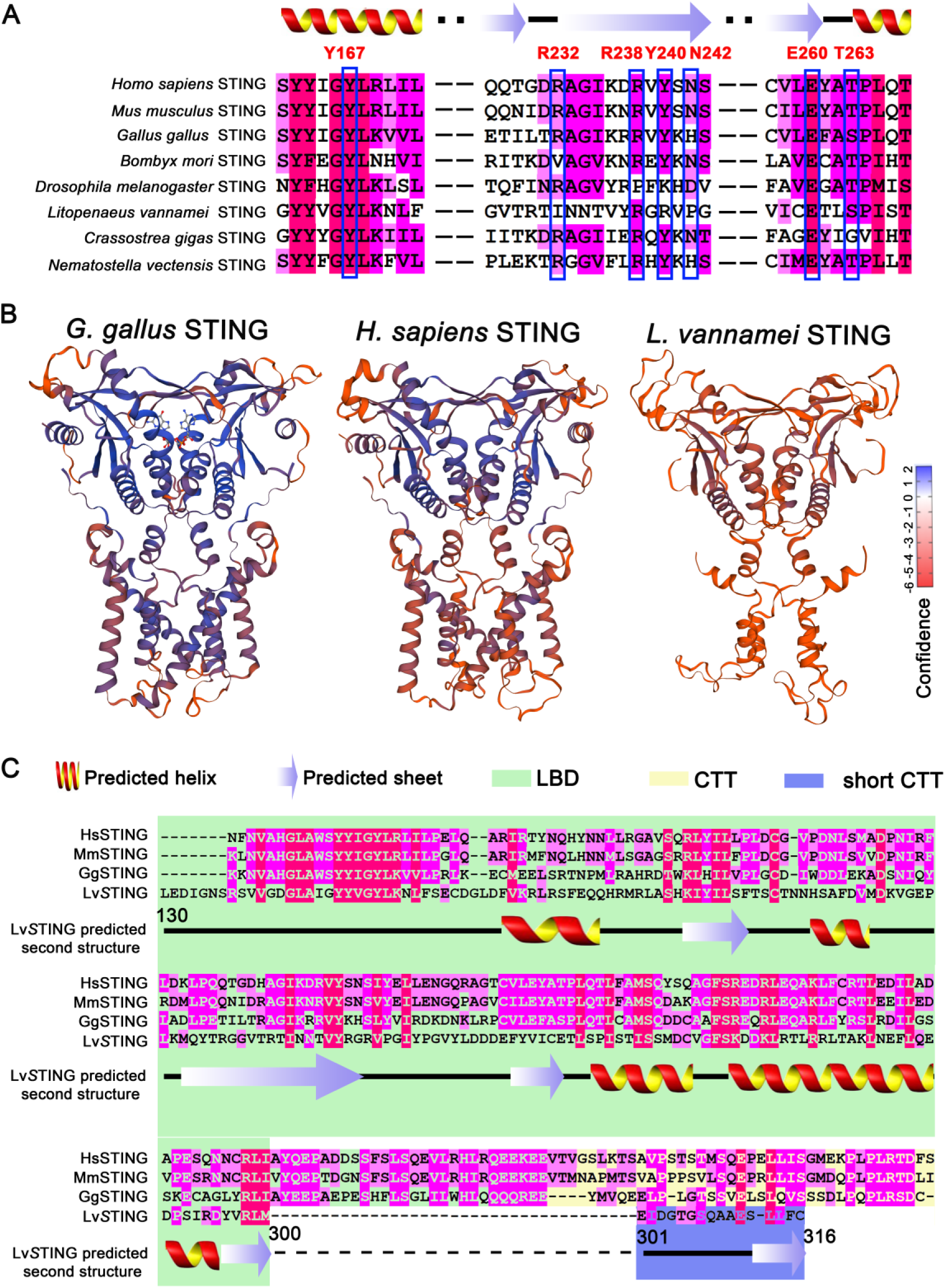
Secondary structure, three-dimensional model, and sequence alignment analysis of shrimp STING. (A) Schematic representation of the architectures of ligand-binding domains (LBDs) from vertebrate and invertebrate STING homologs. The boxed amino residues were crucial to CDN-binding in different STING homologs including *Homo sapiens* STING (HsSTING), *Mus musculus* (MmSTING), *Gallus. gallus* STING (GgSTING), *Bombyx mori* STING, *Drosophila melanogaster* STING, *L. vannamei* STING (LvSTING), *Crassostrea gigas* STING and *Nematostella vectensis* STING. (B) The 3D models of GgSTING, HsSTING and LvSTING predicted by SWISS-Model. (C) Schematic representation of predicted secondary structure of HsSTING, MmSTING, GgSTING and LvSTING. Numbers indicated the locations of amino acids in LvSTING.

Previous study showed that several amino acids at positions Y167, R232, R238, Y240, N242, E260, and T263 in the CDN binding regions of HsSTING are critical to its ligand (2′3′-cGAMP)-binding-induced IFN pathway activation [30]. The overall CDN-binding domains were found to be highly conserved by sequence alignment in several STING homologs, particularly at sites, including Y152, R229, E251, and T254 in LvSTING, which are identical to HsSTING (Figure 1A). Using SWISS-MODEL to predict three-dimensional (3D) structures of STING homodimers based on the template PDB (6nt7.1) from the *Gallus gallus* STING (GgSTING) [31], we observed that LvSTING displayed a homodimeric complex state that is similar to HsSTING and GgSTING (Figure 1B). Sequence alignment on the CTDs of STING homologs showed that LvSTING has helix and sheet arrangements similar to those of HsSTING, *Mus musculus* STING (MmSTING), and GgSTING but has a shorter CTT (sCTT) without the classical pLxIS motif and typical “PLPLRT/SD” motif, which are essential for the recruitment of IRF3 and TBK1 that initiate downstream IFN signaling in vertebrates (Figure 1C). Nevertheless, the above analysis indicated that LvSTING has potential ligand-binding ability for some CDNs.

### LvSTING provides protection against DNA virus infection in shrimps

Given the conservative evolution of LvSTING, we assumed that LvSTING plays a key role in host defense against DNA virus. We purified the expressed protein and prepared a polyclonal anti-LvSTING antibody specific to LvSTING (Figure 2A). After detecting the protein levels of LvSTING in some immune-related tissues and found that LvSTING was abundant in hepatopancreas, intestines, hemocytes, and gills (Figure 2B). The expressional levels of LvSTING were up-regulated in hemocytes and intestines during challenge with two shrimp DNA viruses: WSSV and DIV1 (Figure 2C). These results suggested that LvSTING can be activated by up-regulated expression during DNA virus infection.

**Figure 2.**
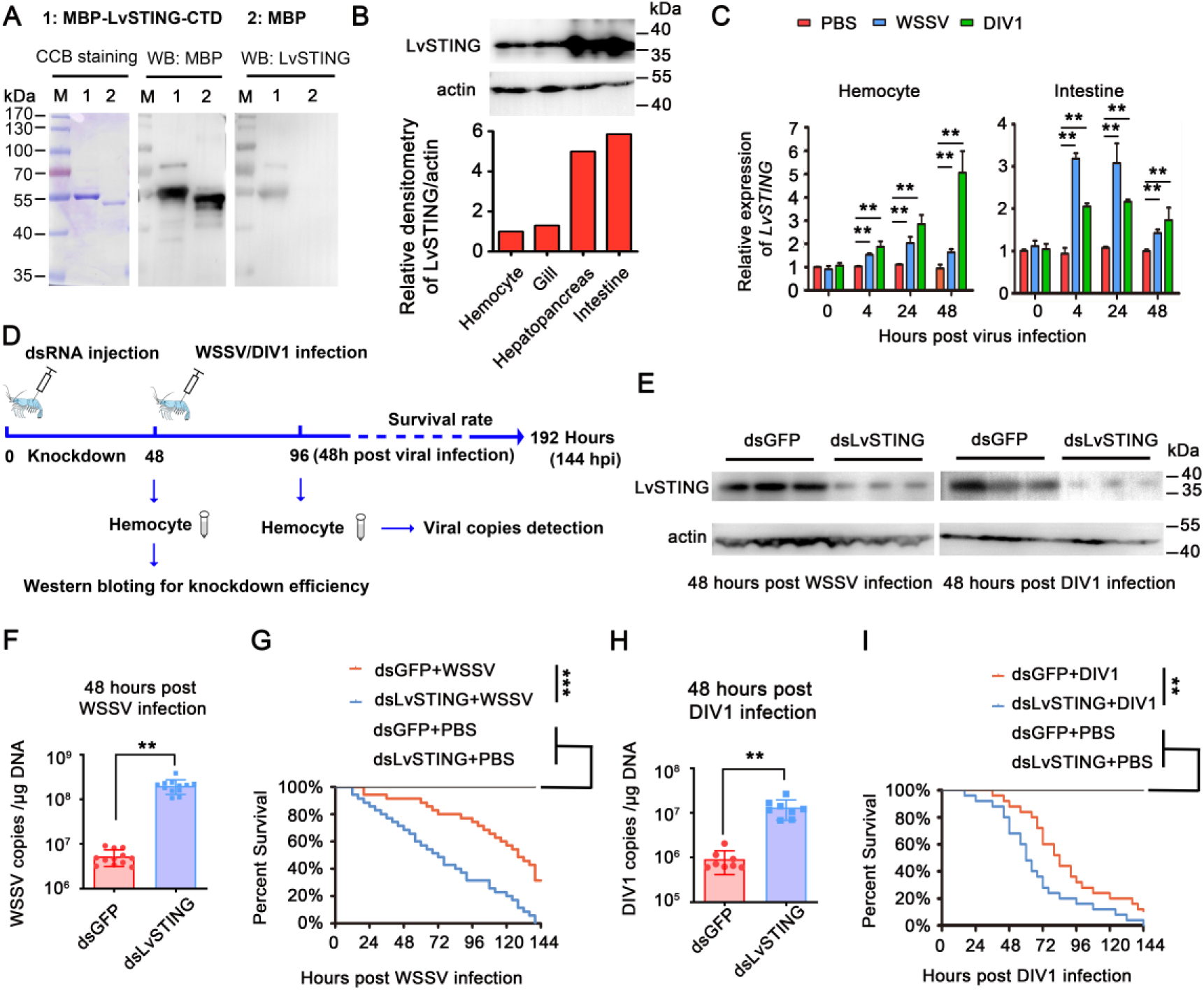
Shrimp STING was required for host defense against DNA viruses. (A) LvSTING antibody was specific to LvSTING detection. Recombinant proteins, including MBP-LvSTING-CTD and MBP, were expressed in Rosseta cells, purified by amylose resins, and detected by CBB staining. Western blotting assays confirmed that the prokaryote-expressed STING protein was recognized by the prepared antibody. (B) The tissue distribution of LvSTING was detected by Western blotting (upper panel). Quantitative analysis of expression levels of LvSTING by ImageJ software corresponded to Western blot results (lower panel). (C) LvSTING expressional patterns responding to the challenges of WSSV, DIV1 and PBS (as a negative control) at different time points in hemocytes and intestine was detected by qRT-PCR. (D) Schematic representation of the procedures for investigating the antiviral function of LvSTING *in vivo*. (E) The RNA interference (RNAi) efficiencies in hemocytes were detected using Western blotting during WSSV or DIV1 challenges. (F, H) Virus copies in hemocytes tissue of dsRNA-LvSTING or dsRNA-GFP (as control) treated shrimp at 48 h post WSSV (F) or DIV1 (H) infection. A Student’s *t*-test was applied (**\*\****p* < 0.01). (G, I) Survival rates of WSSV (G) or DIV1 (I) infected shrimp injected with dsRNA-LvSTING or dsRNA-GFP. Shrimp survival was monitored every 4 h after viral infection. Differences between groups were analyzed with log-rank test using GraphPad Prism 5.0 (**\*\****p* < 0.01, **\*\*\****p* < 0.001). All experiments were replicated two times with similar results.

To investigate the function of LvSTING during DNA virus infection, we knocked down the expression of LvSTING in virus-infected shrimp using RNAi strategy (Figure 2D). The knockdown efficiency was detected in hemocytes via Western blotting, and we observed that protein levels of LvSTING were markedly down-regulated at 48 h after dsRNA injection (Figure 2E). Virus loads in shrimp hemocyte tissues were detected via using absolute quantitative PCR. We observed that the average of viral DNA burden significantly increased after LvSTING was silenced with a 38.58-fold increase 48 h after WSSV infection and 14.51-fold increase 48 h after DIV1 infection (Figure 2F, 2H). Parallel experiments were performed to record the survival rates of LvSTING-silenced shrimp during WSSV or DIV1 infection. Compared with the dsRNA–GFP control group, the dsRNA-LvSTING-treated group showed significant decrease in survival rate during WSSV infection (χ^2^: 22.32, *p* < 0.0001) and during DIV1 infection (χ^2^: 11.44, *p* = 0.0096 < 0.001), respectively (Figure 2G, 2I). Taken together, STING is essential to defense against DNA viruses, such as WSSV and DIV1, in shrimp.

### LvSTING binds to several CDN species

Given the conservation of the LBD in LvSTING, we performed pull-down studies to explore whether LvSTING can bind to CDN species, such as c-di-GMP, c-di-AMP, 3′3′-cGAMP and 2′3′-cGAMP. HsSTING were tested as a positive control. We observed that HsSTING and LvSTING were able to bind to c-di-GMP, c-di-AMP, 3′3′-cGAMP and 2′3′-cGAMP (Figure 3A), but did not interact with poly (I:C) (Figure 3–figure supplement 1) that excluded the possible interaction between STING and CDNs attributed to a non-specific and electrostatic action. Next, we determined whether the LBD domain of LvSTING is responsible for CDN binding. Several plasmids containing the sequences of LvSTING truncating forms were constructed and expressed in *Drosophila* S2 cells (Figure 3B). Some pull-down experiments were then carried out using 2′3′-cGAMP-coupled agarose and detected by Western blotting with anti-HA antibody. The results showed that the peptides containing the 130–300 residues of LvSTING are fully capable of binding to 2′3′-cGAMP (Figure 3C).

**Figure 3.**
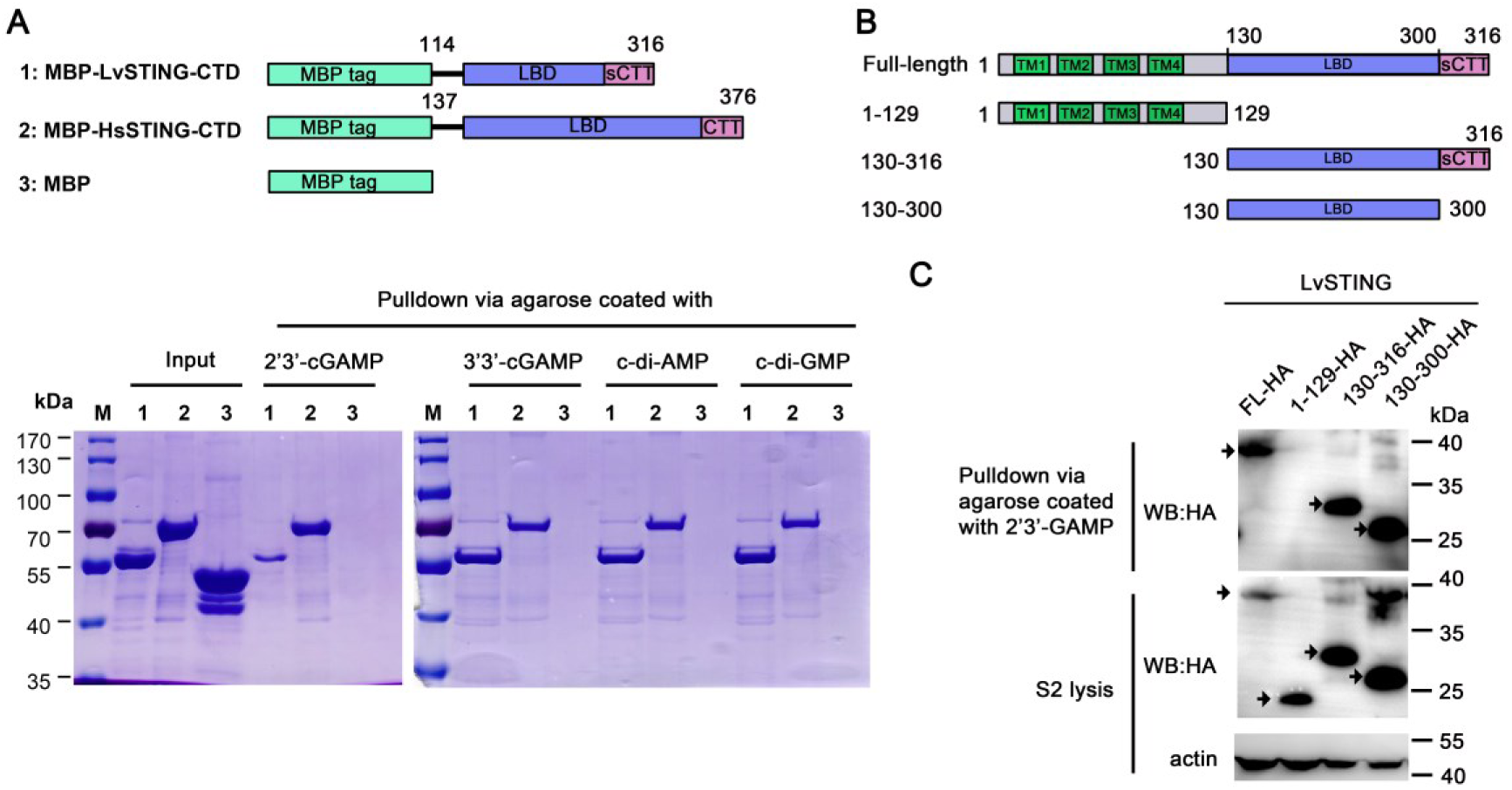
Shrimp STING could bind to several CDNs including 2′3′-cGAMP. (A) Pull-down assay was used to determine the interactions between STING proteins and CDNs. Plasmids, including MBP-HsSTING-CTD, MBP-LvSTING-CTD (positive control), and MBP (negative control) were constructed for pull-down assays (upper panel). Pull-down assay confirmed that MBP-LvSTING-CTD and MBP-HsSTING-CTD interacted with several CDNs including 2′3′-cGAMP, 3′3′-cGAMP, c-di-AMP and c-di-GMP (lower panel). (B) Schematic representations of various LvSTING deletion mutants. (C) The ligand binding domain (LBD) of LvSTING (region 130–300) bound to 2′3′-cGAMP. Various fragments of LvSTING expressed in *Drosophila* S2 cells were pulled down with agarose coated with 2′3′-cGAMP. All experiments were replicated three times with similar results.

### Shrimp STING combines with IKKε and IRF to establish a functional signaling axis that induces Vago4 activation in *Drosophila* S2 cells

To understand the molecular mechanism underlying LvSTING-mediated antiviral immunity, we determined whether LvSTING can induce the activation of the IRF–Vago antiviral axis, which is parallel to the IRF–IFN antiviral axis in mammals [28]. A series of various plasmids coding different regions of LvSTING, LvIRF (homologous to mammalian IRF3) [28], and LvIKKε (homologous to mammalian TBK1/IKKε) [32] were constructed (Figure 4A). Co-immunoprecipitation (Co-IP) was conducted in *Drosophila* S2 cells to determine whether LvSTING can interact with LvIRF and LvSTING. Co-IP combined with Western blotting assays showed that the sCTT (301–316 amino acids) of LvSTING facilitated its interaction with LvIRF and LvIKKε (Figure 4B, 4D). The C-terminal region (120–362 amino acids) of LvIRF and N-terminal region (1–306 amino acids) of LvIKKε contributed to interacting with LvSTING (Figure 4C, 4E). LvIRF induces the expression of LvVago4 (an interferon functional analog) by directly binding to its promoter [28]. Next, we explored whether LvSTING, LvIRF and LvIKKε can form a functional signaling axis to activate the promoter activities of *LvVago4*. Given the absence of immortalized cell line in shrimp, *Drosophila* S2 cells are alternatives for detecting the effect of the LvSTING–LvIKKε– LvIRF axis on LvVago4 induction. Besides, no IRF homolog coding sequence has been discovered in *Drosophila* genome, which can avoid the interference from endogenous IRF protein [33]. We observed that ectopically co-expressed proteins, including LvIKKε and LvIRF, and the full-length of LvSTING, except the 1–300 amino acids of LvSTING, strongly induced the promoter activities of *LvVago4* with or without 2′3′-cGAMP treatment (Figure 4F). Accordingly, the full-length of LvSTING, but not the sCTT deleted mutant of LvSTING, can induce *LvVago4* in a concentration dependent manner in *Drosophila* S2 cells (Figure 4–figure supplement 1). These results indicated that the sCTT (301–316 amino acids) of LvSTING is required in the formation of the functional complex of LvIKKε, LvIRF and LvSTING for the initiation of downstream signaling.

**Figure 4.**
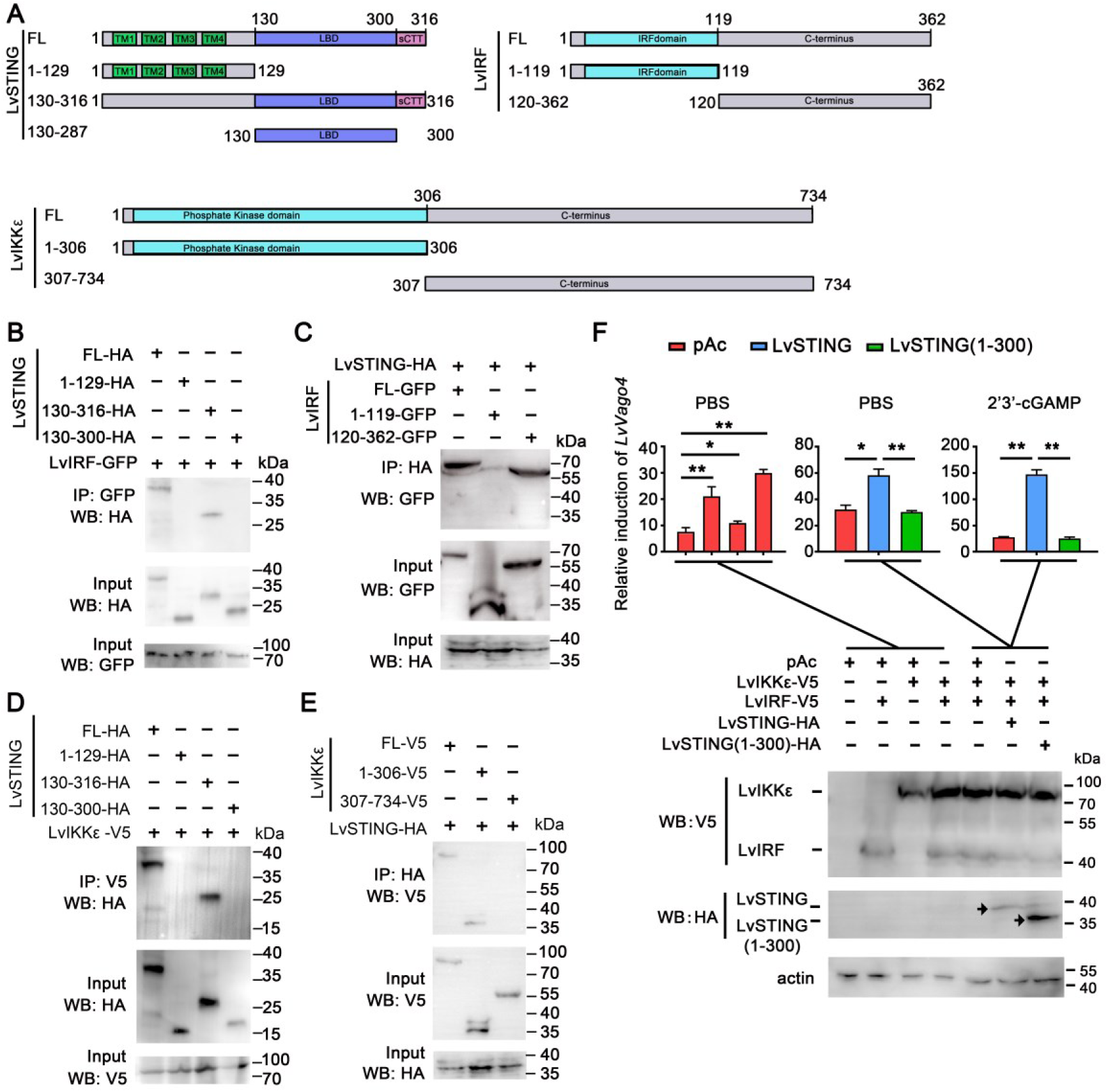
Shrimp STING–IKKε–IRF complex induced the promoter activities of *LvVago4* in cells. (A) Schematic representations of various deletion mutants of LvSTING, LvIRF and LvIKKε. (B) Interactions between GFP-tagged LvIRF and HA-tagged full-length or truncated LvSTING were analyzed by Co-IP assays. (C) Interactions between HA-tagged LvSTING and GFP-tagged full-length or truncated LvIRF were analyzed by Co-IP assays. (D) Interactions between V5-tagged LvIKKε and HA-tagged full-length or truncated LvSTING were analyzed by Co-IP assays. (E) Interactions between HA-tagged LvSTING and V5-tagged full-length and truncated LvIKKε was analyzed by Co-IP assays. (F) Relative induction of the promoter activities of *LvVago4* mediated by LvIRF-V5, LvIKKε-V5 and LvSTING-HA or LvSTING-1– 300-HA in *Drosophila* S2 cells treated by 2′3′-cGAMP using dual luciferase assays. Ectopically expressed proteins were detected by Western blotting. Actin was used as a protein loading control. The bars indicated the mean ± SD of the luciferase activities (*n* = 3). The data was analyzed statistically by student’s *t*-test (**\*\****p* < 0.01). All experiments were replicated three times with similar results.

### Exogenous DNA triggers the activation of STING–IRF–Vago4 signaling axis in shrimp hemocytes

In mammals, exogenous DNA can induce the activation of a STING–IRF3–IFN cascade [34]. As mentioned above, shrimp STING can bind to the several second messengers of the cGAS–STING pathway (Figure 3) and can combine with IKKε and IRF to form a functional signaling axis that induces Vago4 activation in *Drosophila* S2 cells (Figure 4). Thus, we explored whether exogenous DNA can activate the STING– IRF–Vago4 signaling axis in shrimp cells. In the primary cultured shrimp hemocyte cells, we observed that the transfection of exogenous DNA (dsDNA1 and dsDNA2, whose sequences were selected from shrimp viral genomes of WSSV and DIV1 respectively, showed in Supplementary Table S2) can induce LvIRF dimerization 4 h after stimulation, as detected by Western blotting with anti-LvIRF antibody in Native PAGE (Figure 5A). The immunofluorescence assay demonstrated the elevated nuclear translocation levels of LvIRF stimulated by dsDNA1/2 (Figure 5B). Meanwhile, the expression levels of LvVago4 were strongly induced by DNA transfection in shrimp hemocytes (Figure 5C). These results suggested that the transfection of exogenous DNA can activate LvIRF–LvVago4 cascade in primary cultured shrimp hemocyte cells. Next, we explored whether LvSTING is crucial for LvIRF–LvVago4 activation by exogenous DNA transfection. Hemocytes from LvSTING-silenced shrimps were harvested and planted in cell culture dishes. After transfection with dsDNA1 or dsDNA2, we observed that LvIRF dimerization was dramatically impaired in the LvSTING-silenced shrimp hemocytes (Figure 5D). PBS and dsGFP treatments were used as controls. The knockdown of LvSTING significantly suppressed dsDNA1/2-induced LvIRF nuclear translocation (Figure 5E-F) and the expression of LvVago4 (Figure 5G). The data demonstrated that LvSTING is required for dsDNA-induced IRF–Vago cascade activation in shrimp hemocytes.

**Figure 5.**
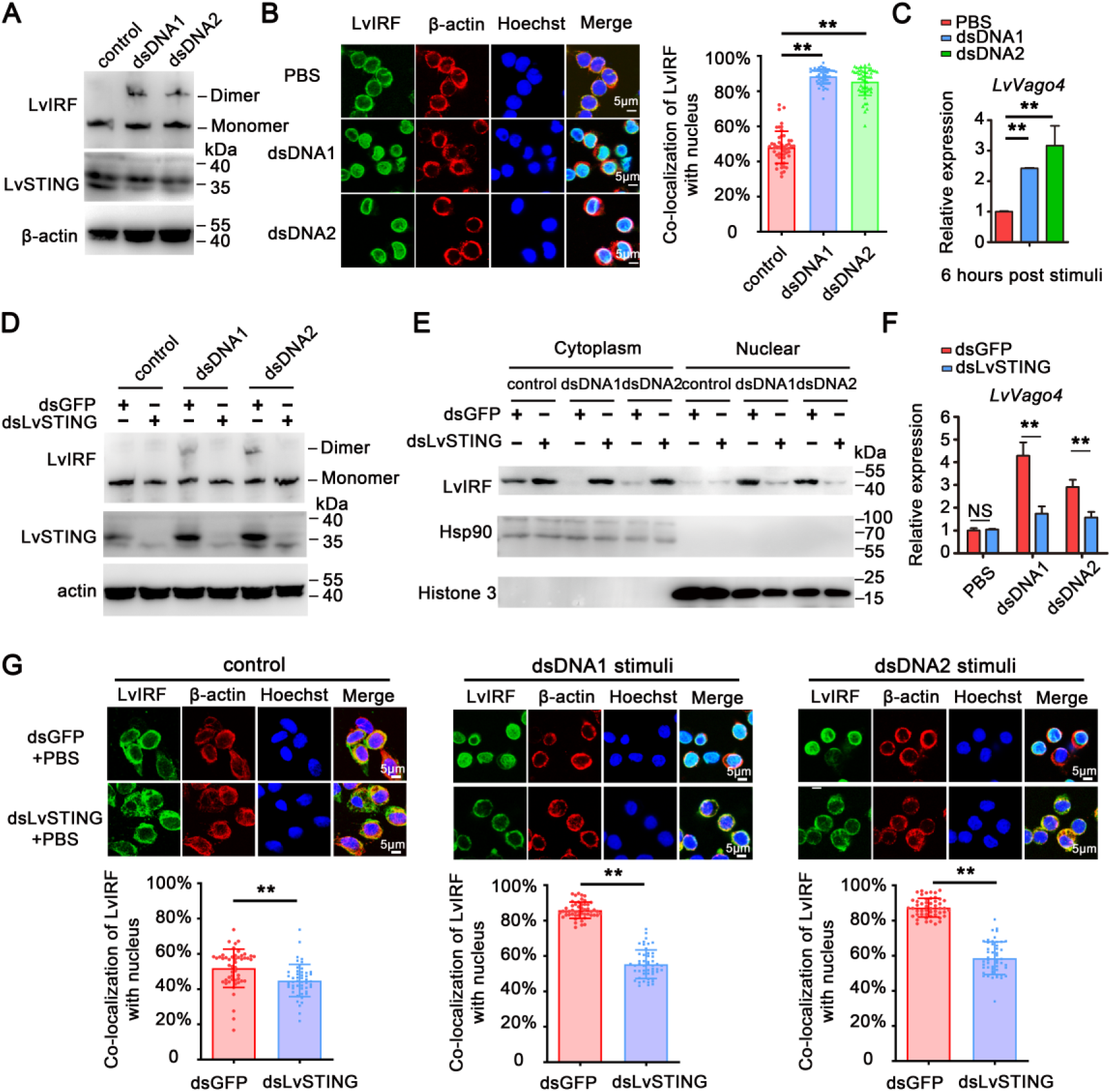
Exogenous dsDNA activated shrimp STING–IRF–Vago4 axis in cells and *in vivo*. (A) Dimerization stimulation of LvIRF in primary culture shrimp hemocytes transfected with exogenous dsDNA1/2. (B) Exogenous dsDNA1/2 transfection induced LvIRF nuclear translocation in primary culture shrimp hemocytes. Statistical analysis of the co-localization of LvIRF and nucleus in hemocytes (*n* = 50 cells) corresponding to immunofluorescence (right panel). (C) Exogenous dsDNA1/2 transfection activated the transcriptional expression of *LvVago4* in primary culture shrimp hemocytes. (D-G) Silencing of LvSTING *in vivo* impaired exogenous dsDNA1/2 mediated LvIRF dimerization stimulation (D), nuclear translocation (E, G), and *LvVago4* expression (F). Statistical analysis of co-localization of LvIRF and nucleus in hemocytes (*n* = 50 cells) corresponding to immunofluorescence (lower panel of G). Data from (C) and (F) were analyzed statistically by student’s *t*-test (NS, not significant, **\*\****p* < 0.01). Data from (B) and (G) (*n* = 50 cells) were calculated by ImageJ software and analyzed statistically by student’s *t*-test (**\****p* < 0.05, **\*\****p* < 0.01). All experiments were replicated three times with similar results.

### Shrimp cGAS homolog is not involved in the DNA-intrigued activation of IRF– Vago4 signaling axis in hemocytes

Mammalian cGAS homologs can directly sense cytosolic DNA and produce 2′3′-cGAMP to activate the type I interferon pathway [6]. A cGAS homolog named LvcGAS (also named Mab21-containing protein, LvMab21cp) was cloned and identified in *L. vannamei* [35]. Thus, we determined whether LvcGAS plays a role in parallel interferon-like (IRF–Vago) antiviral signaling axis responding to DNA virus infection or DNA stimulation. First, the silencing of LvcGAS alone decreased LvVago4 expression in the absence of WSSV infection (Figure 6–figure supplement 1A), and LvVago4 expression was still down-regulated in LvcGAS-silenced shrimp after WSSV infection (Figure 6–figure supplement 1B). Then, we observed that shrimps with knocked down LvcGAS expression had elevated viral loads and were more susceptible to WSSV infection (Figure 6–figure supplement 1C-D). These results strongly suggested that LvcGAS played key roles in host defense against WSSV infection and in the regulation of LvVago expression *in vivo*. We observed that the silencing of LvcGAS substantially attenuated the dimerization and nuclear translocation of IRF in response to WSSV infection. The expression of LvVago (Figure 6A), dimerization (Figure 6B), and nuclear translocation (Figure 6B) of IRF were not affected in hemocytes from LvcGAS-silenced shrimps in response to exogenous DNA stimulation. Vertebrate cGAS contains a unique zinc ribbon domain required for DNA sensing [36], whereas the zinc-ribbon domain is not present in LvcGAS [35]. Indeed, the pull-down experiment showed that LvcGAS did not bind to dsDNA (Figure 6D) in contrast to human cGAS. Taken together, these results suggested that shrimp LvcGAS is involved in the WSSV, but not DNA, intrigued activation of the IRF–Vago4 signaling axis.

**Figure 6.**
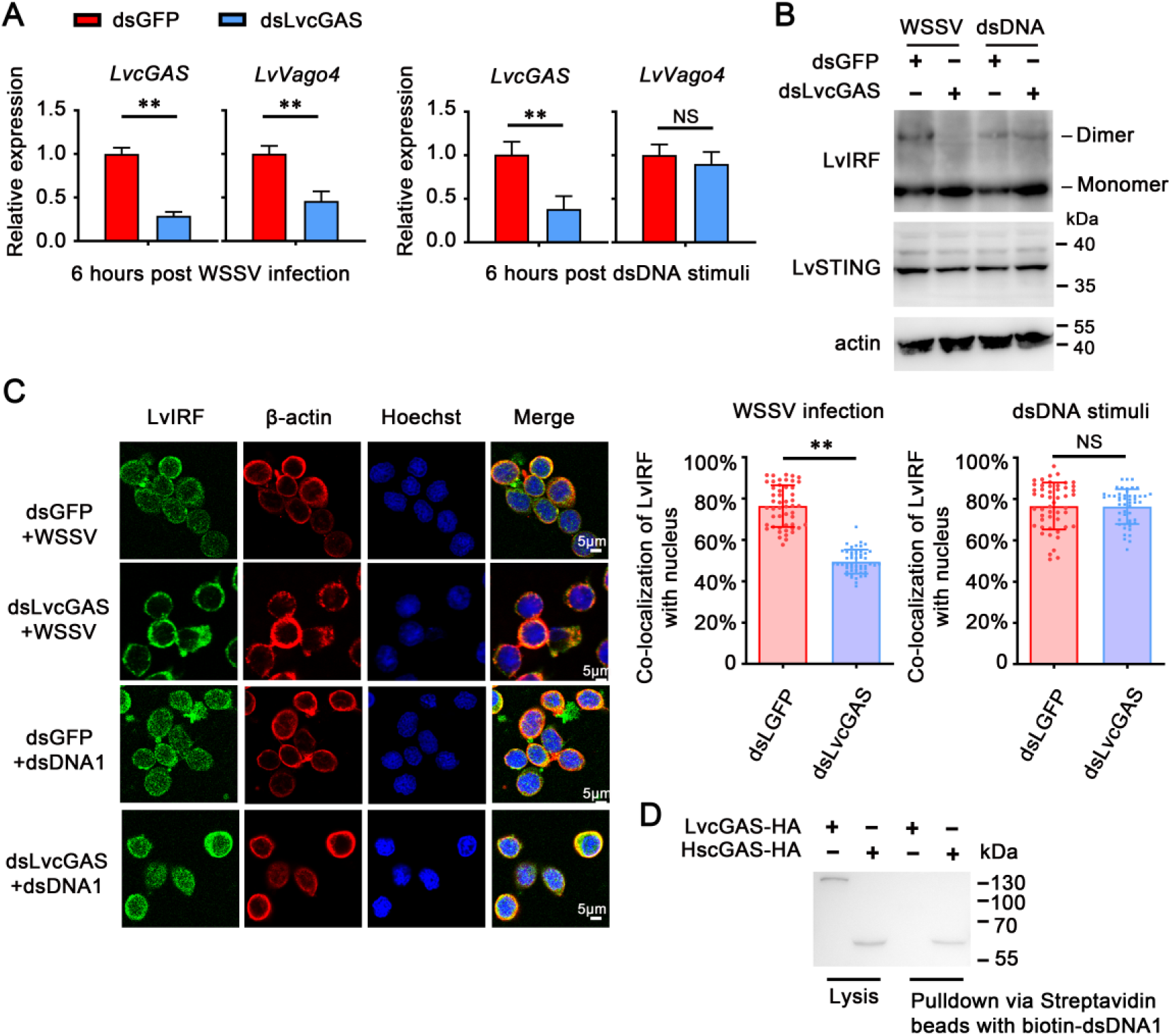
Shrimp cGAS homolog might not participate in exogenous dsDNA-intrigued STING–IKKε-IRF–Vago4 axis in cells and *in vivo.* (A) *LvVago4* transcription expression in WSSV-infected (left panel) or dsDNA-transfected (right panel) primary culture hemocytes from dsRNA-LvcGAS-treated shrimps. (B) Dimerization stimulation of LvIRF in WSSV-infected or dsDNA-transfected primary culture hemocytes from dsRNA-LvcGAS-treated shrimps. (C) WSSV infection or exogenous dsDNA transfection induced LvIRF nuclear translocation in primary culture shrimp hemocytes from dsRNA-LvcGAS-treated shrimps. Statistical analysis of co-localization of LvIRF and nucleus in hemocytes (*n* = 50 cells) corresponding to immunofluorescence (right panel). (D) LvcGAS did not bind to dsDNA in pull-down assay. LvcGAS-HA and HscGAS-HA expressing in *Drosophila* S2 cells were pulled down with agarose coated with biotin-dsDNA1. All experiments were replicated three times with similar results.

### Shrimp STING binds to cytosolic dsDNA in cells

Given that LvcGAS is not responsible for activating IRF–Vago axis in response to exogenous dsDNA stimulation, we determined whether LvSTING can bind to dsDNA. We performed pull-down studies with biotin-dsDNA1, biotin-dsDNA2, and biotin-poly (I:C). The results showed that the peptide spanning amino acid residues 130–300 of LvSTING can bind to dsDNA fragments rather than to dsRNA mimics (Figure 7A). Pull-down assays by streptavidin beads and Western blotting by anti-LvSTING antibody showed that endogenous LvSTING from hemocytes can bind to dsDNA (Figure 7B). Competition experiments indicated that unbiotinylated dsDNA can effectively compete with biotinylated dsDNA for LvSTING binding to hemocytes (Figure 7B). We then performed pull-down studies to explore which regions of LvSTING LBD are essential for dsDNA binding by expressing STING fraction deletion variants (LvSTINGΔ145–184, LvSTINGΔ185–224, and LvSTINGΔ225–264). LvSTINGΔ225–264 were observed to be less pulled down when biotinylated dsDNA1/2 was used, whereas LvSTINGΔ145-184, LvSTINGΔ185–224, and LvSTINGΔ225–264 weakened ability to interact with 2′3′-cGAMP (Figure 7C). To explore whether the induced promoter activities of *LvVago4* was required for dsDNA binding by LvSTING, dual-luciferase reporter assays were conducted with plasmids containing wild type or fraction deletion (LvSTINGΔ145–184, LvSTINGΔ185–224, and LvSTINGΔ225–264) of LvSTING. We found that LvIKKε and LvIRF together with LvSTING wild type, but not these delete mutants, can form a functional 2′3′-cGAMP-intrigued STING–IRF–Vago4 signaling axis, and 225–264 deletion of LvSTING can suppress dsDNA-triggered *LvVago4* promoter activities most significantly (Figure 7D). To further define the binding ability of LvSTING in cytosolic dsDNA sensing, the CTD of LvSTING was each cloned into prokaryotic expressing plasmids with maltose-binding-protein (MBP) tags and purified by amylose resin (Figure 7E). The dissociation constant (*Kd*) was determined by MST *in vitro*. Recombinant MBP (rMBP) was used here as a negative control for dsDNA binding. As Figure 7E shows, the binding affinity of LvSTING with dsDNA was 9.21 ± 1.49 μM. Collectively, these data suggested that the 225–264 amino acids contribute to the binding activity of LvSTING and the DNA binding ability of LvSTING is crucial for inducing downstream signaling.

**Figure 7.**
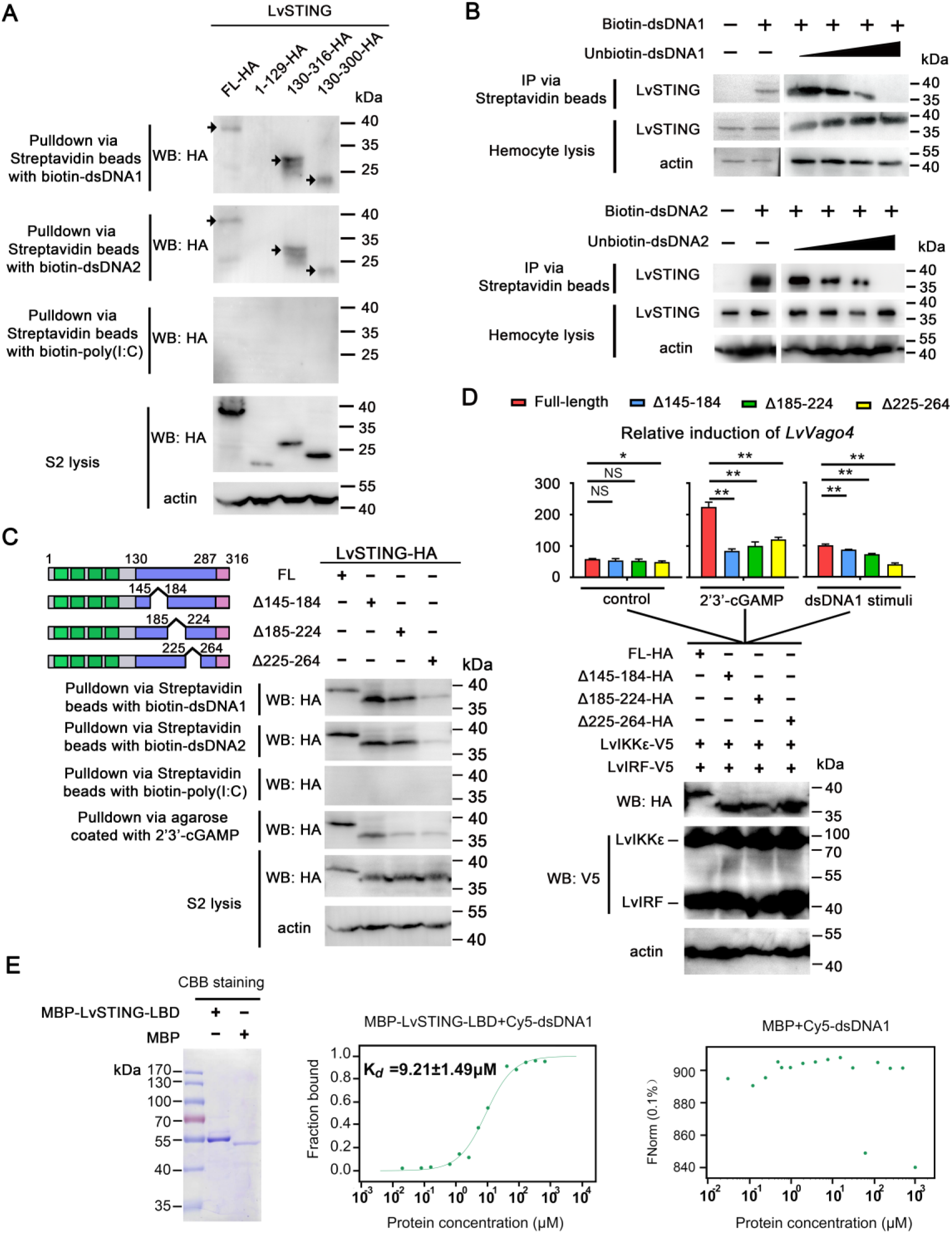
Shrimp STING bound to dsDNA. (A) Pull-down analysis of various LvSTING deletion mutants binding to dsDNA *in vitro*. Various fragments of LvSTING expressed in *Drosophila* S2 cells were pulled down with agarose coated with biotin-dsDNA or biotin-poly (I:C). (B) Endogenous LvSTING interacted with dsDNA. LvSTING in shrimp hemocytes was precipitated by biotin-dsDNA1/2, and detected using Western blotting with anti-LvSTING antibody. (C) Pull-down assay of various LvSTING LBD-deletion mutants binding to dsDNA *in vitro*. *Drosophila* S2 cells were transfected with LvSTING full length-HA, LvSTINGΔ145–184-HA, LvSTINGΔ185– 224-HA, and LvSTINGΔ225–264-HA for 48 h. Cell lysis was precipitated with streptavidin beads or agarose and analyzed by immunoblotting. (D) Relative induction of the promoter activities of *LvVago4* mediated by LvIRF, LvIKKε, LvSTING-WT-HA, LvSTINGΔ145–184-HA, LvSTINGΔ185–224-HA or LvSTINGΔ225–264-HA in *Drosophila* S2 cell treated with 2′3′-cGAMP or dsDNA1 using dual luciferase assays. Ectopic expressed proteins were showed by Western blotting. Actin was used as a protein loading control. The bars indicated the mean ± SD of the luciferase activities (*n* = 3). Data were analyzed statistically by student’s *t*-test (**\*\****p* < 0.01). (E) Recombinant proteins, including MBP-LvSTING-CTD, and MBP, were expressed in Rosseta cells and purified by amylose resin (left panel). The affinity of MBP-LvSTING-CTD (middle panel) or MBP (right panel) binding to Cy5-dsDNA1 was measured by MST. MST values were normalized to the fraction bound and dissociation constant (*Kd*) of each group. All experiments were replicated three times with similar results.

### Shrimp STING facilitates viral DNA sensing to induce the activation of IRF– Vago4 antiviral axis *in vivo*

We demonstrated that shrimp STING can facilitate synthetic DNA sensing and is essential for defense against DNA virus infection. This result prompted us to determine whether LvSTING facilitates host immune system in sensing viral DNA during infection. Thus, we performed chromatin immunoprecipitation (ChIP) experiments were performed to detect the probability of LvSTING in recognizing viral DNA *in vivo*. The hemocytes from WSSV-infected shrimps 48 h after infection were harvested for ChIP assay with anti-LvSTING antibody. PCR was used in detecting ChIP products with several pairs of primers, which were designed for targeting the random regions of the WSSV genome including the two repeated sequence regions of Hr1 and Hr5, five promoter regions of *wsv069*, *wsv078*, *wsv079*, *wsv249*, and *wsv403*, and three coding sequence regions of *wsv184*, *VP15*, and *VP28* (Figure 8A). We observed that all these regions could be immunoprecipitated by anti-LvSTING antibody, but not the IgG control (Figure 8A). These results suggested that endogenous LvSTING can bind to viral DNA.

**Figure 8.**
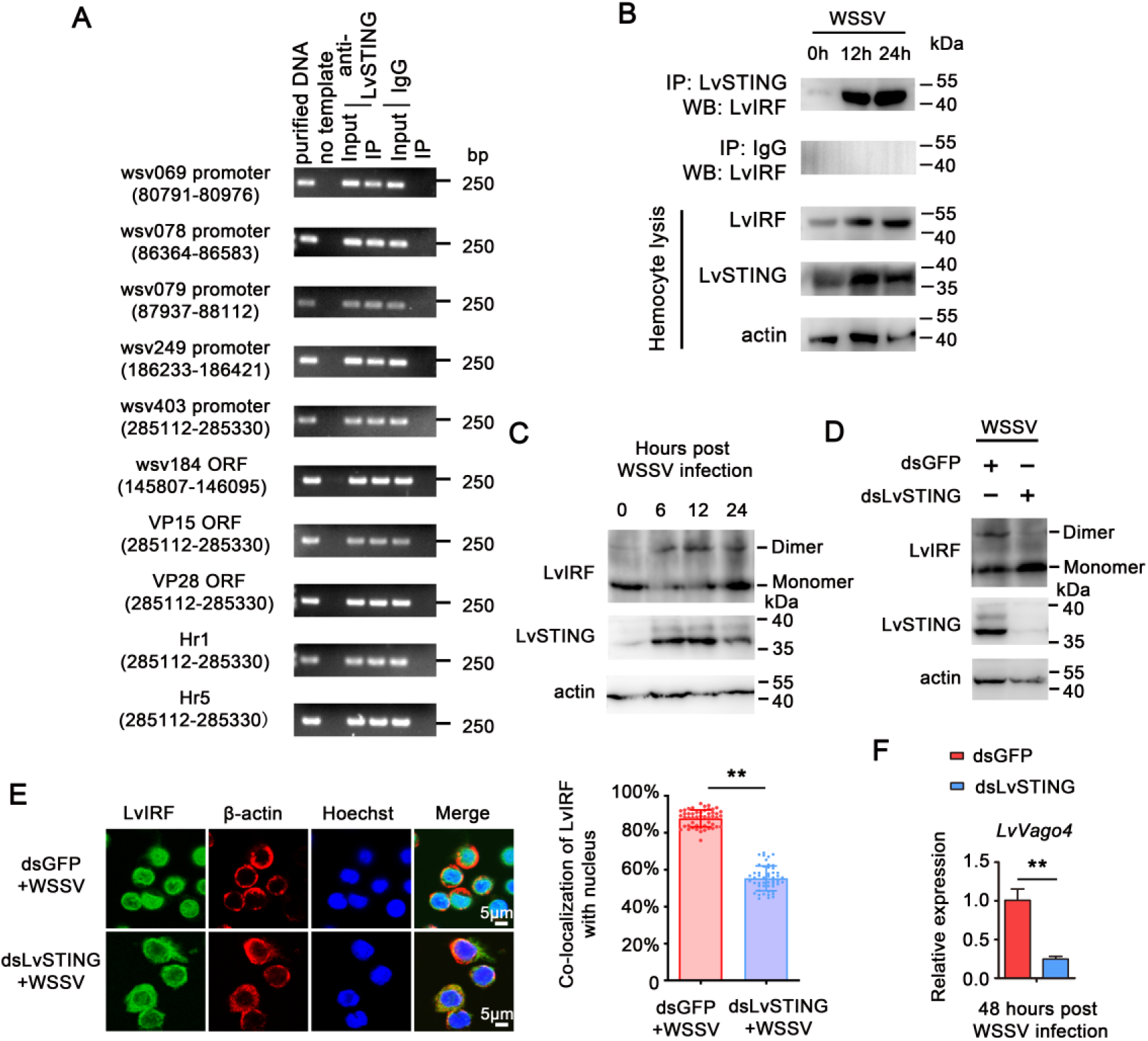
Shrimp STING directly recognized viral DNA during infection to activate IRF–Vago4 axis *in vivo*. (A) Multiple regions of WSSV genome were sensed by endogenous LvSTING during infection. ChIP assays were performed in WSSV-infected hemocytes by the anti-LvSTING antibody or IgG (as negative control). Multiple regions, including two highly repeated sequences (Hr1, and Hr5), five promoter regions (*wsv069*, *wsv078*, *wsv079*, *wsv249 and wsv403*) and three coding regions (*wsv184*, *VP15*, and *VP28*) bound to endogenous LvSTING. Five percent of the samples before immunoprecipitation were detected as input controls. (B) Increased interaction levels of LvSTING–LvIRF were observed 0, 12, and 24 h after WSSV infection *in vivo*. (C) Increased IRF dimerization at 0, 12, and 24 h after WSSV infection *in vivo*. (D–F) Silencing of LvSTING weakened LvIRF dimerization levels (D), nuclear translocation (E), and the transcriptional expression of *LvVago4* (F) in response to WSSV infection. Hemocytes were collected 48 h after WSSV infection, and then subjected to Western blot analysis, immunofluorescence staining, or qRT-PCR. Co-localization of LvIRF and nucleus in hemocytes (*n* = 50 cells) was calculated by ImageJ and analyzed statistically by student’s *t*-test (**\*\****p* < 0.01). The data (F) were analyzed statistically by student’s *t*-test (**\*\****p* < 0.01). All experiments were replicated three times with similar results.

To further confirm whether LvSTING assistance to viral DNA sensing during WSSV infection can activate LvIRF-dependent innate immunity *in vivo*, we detected a series of activation events downstream of LvSTING. We found that WSSV infection can significantly induce the interactions of LvSTING and LvIRF by endogenous IP analysis (Figure 8B). LvIRF dimerization can be triggered by WSSV infection (Figure 8C). The knockdown of LvSTING, rather than dsGFP control, suppressed WSSV-induced LvIRF dimerization (Figure 8D), LvIRF nuclear translocation (Figure 8E), and LvVago4 expression levels (Figure 8F). Similar results were observed during DIV1 infection (Figure 8–figure supplement 1). Taken together, these data suggested that shrimp STING can facilitate the host immune system sin sensing viral DNA during infection and induces the activation of IRF–Vago4 antiviral axis *in vivo*.

## DISCUSSION

Genetic evidence has shown that cGAS–STING signaling is essential to the activation of type I IFN response by cytosolic DNA in mammals [33]. Given the indispensability of cGAS–STING signaling in mammals, exploring its functionally evolutionary origins is necessary. Scientists have devoted considerable effort to identifying the orthologs of cGAS and STING in lower organisms, including invertebrates and even bacteria. Human cGAS is composed of an unstructured and poorly conserved N terminus (amino acid residues 1–160) and a highly conserved C terminus (160–513). The positively charged N-terminal fragment plays a critical role in DNA binding [20]. Most cGAS homologs in vertebrates and a cephalochordate (*Branchiostoma floridae*) have long N-terminal fragments with an average length of 167 amino acids. By contrast, invertebrate cGAS-like homologs contain extremely short N-terminal fragments, such as ∼70 amino acids in *Nematostella vectensis* [26]. Similarly, human Mab21-like proteins, lacking the ability to detect DNA, contain extremely short N-terminal tails [26]. These features strongly suggest that the 167-amino-acid-long N-terminal tail has evolved in chordate or vertebrate lineage and seems to be a cGAS-specific adaptation. Moreover, the C-terminal fragment of human cGAS contains two conserved domains, nucleotidyltransferase (NTase) core domain, and Mab21 domain [6]. A zinc ribbon structural domain is inserted into the Mab21 region and is essential for a metal coordination and interaction with the groove of DNA, which might endow the C-terminal fragment as a molecular “ruler” for scaling the specificity of cGAS toward DNA [37]. Although some cGAS-like homologs from choanoflagellates, cnidarians, insects, and cephalochordate show conservation on the NTase core and Mab21 domains, the absence of a zinc ribbon domain can make most invertebrate cGAS-like homologs lose their ability to sense DNA [26]. In fact, the *L. vannamei* cGAS-like homolog contains a short N-terminus and a Mab21 domain but does not possess zinc ribbons [35]. Hence, LvcGAS cannot directly interact with DNA (Figure 6D). Notably, the ectopic expression of LvcGAS cannot enhance the dsDNA-activated and LvSTING-dependent production of IFN-β and IFN-ω in HEK293T cells [35], and our results showed a similar result, which demonstrated that LvcGAS did not trigger the dsDNA-intrigued STING–IKKε–IRF–Vago axis *in vivo*. Therefore, whether shrimp cGAS-like proteins participate in innate immune in response to DNA needs to be carefully investigated.

The absence of a zinc ribbon domain in invertebrate cGAS proteins does not mean the absence of DNA sensing-dependent immune pathways in invertebrates. Some reports have shown that DNA can induce potent innate immune responses in invertebrates [25, 38]. In humans, cGAS is the cytoplasmic dsDNA sensor that catalyzes the synthesis of the CDN of 2′3′-cGAMP, which is sensed by STING that induces the expression of type I IFN through the TBK1–IRF3 signaling axis [19, 39–41]. STING can recognize bacterium-derived CDNs, including 3′3′-cGAMP, c-di-GMP, and c-di-AMP [42–44]. Interestingly, STING facilitates dsDNA sensing by directly interacting with DNA sensors that can instigate cytoplasmic DNA-mediated cellular defense responses [16–18]. In *Penaeus monodon*, the DDX41 homolog has been identified as a potential cytosolic DNA sensor that can activate the IFN pathway by interacting with the ectopically expressed mouse STING in HEK293T cells [38]. In this study, we found that LvcGAS is essential for host defense against WSSV infection. Interestingly, LvcGAS is involved in WSSV infection, but not dsDNA, induced the activation of the IRF–Vago axis. These findings indicated that other proteins might participate in interactions between LvcGAS and the downstream IRF–Vago axis during a viral infection. The mechanism by which DNA virus infection activates the cGAS mediated interferon-like antiviral response in shrimp requires further study.

The cGAS–STING-mediated antiviral nucleotide signaling is an ancient mechanism against infection in diversity organisms (Figure 9). Animal cGAS-mediated innate immune signaling shows remarkable homology with the components of cyclic oligonucleotide-based antiphage signaling system (CBASS) and clustered regularly interspaced short palindromic repeats (CRISPR) antiphage immunity in diverse prokaryotes (bacteria and archaea) [45]. In CBASS, a bacterial CBASS operon encodes a cGAS/DncV-like nucleotidyltransferases that synthesizes a nucleotide second messenger in response to phage infection [46]. In type III CRISPR, the sensing of phage infection leads to the activation of the protein Cas10, which synthesizes some cyclic oligoadenylate rings that are 3–6 nucleotides in size [47, 48]. These nucleotide second messengers are recognized by related receptors, which in turn activate downstream effectors that execute cell death to destroy host cells and block phage propagation [49–54]. The origin of STING can be traced back to the prokaryote *Flavobacteriaceae* and the choanoflagellate *Monosiga brevicollis* [55], whereas IKKε, TBK1, and IRF are absent in these organisms. The IKKε and TBK1 families arose in early metazoans, such as the poriferan *Amphimedon queenslandica*, nematode *Caenorhabditis elegans*, and insect *D. melanogaster*, but they lack IRF members. The IRF families have been reported in some invertebrates, such as the mollusk *C. gigas* [56] and crustacean *L. vannamei* [28]. Therefore, whether a functional STING–TBK1/IKKε–IRF signaling axis arise in invertebrates is an interesting topic. In vertebrates, this signaling axis requires that STING has a functional C-terminal tail (CTT), which is indispensable for recruiting TBK1/IKKε and IRF3 and for downstream transducing signals [21]. Previous assay indicates that invertebrate STING lacks the functional CTT, suggesting there may not be STING–TBK1/IKKε–IRF signaling axis in invertebrates [26]. In our study, we found that shrimp STING lacks the conserved PLPRT/SD motif whereas having a sCTT domain within a β-sheet (Figure 2C). Nonetheless, the sCTT domain of shrimp STING is involved in mediating interactions with shrimp IKKε and IRF and establishes a regulatory cascade to induce antiviral factor LvVago4 *in vitro* and *in vivo*. Functionally, STING is essential to activate downstream IKKε–IRF–Vago antiviral signaling axis in response to exogenous DNA or DNA virus. Taken together, these results strongly suggested that shrimp indeed has a functional STING–IKKε–IRF–Vago antiviral signaling axis parallel to the STING–TBK1–IRF–IFN signaling cascade in mammals.

**Figure 9.**
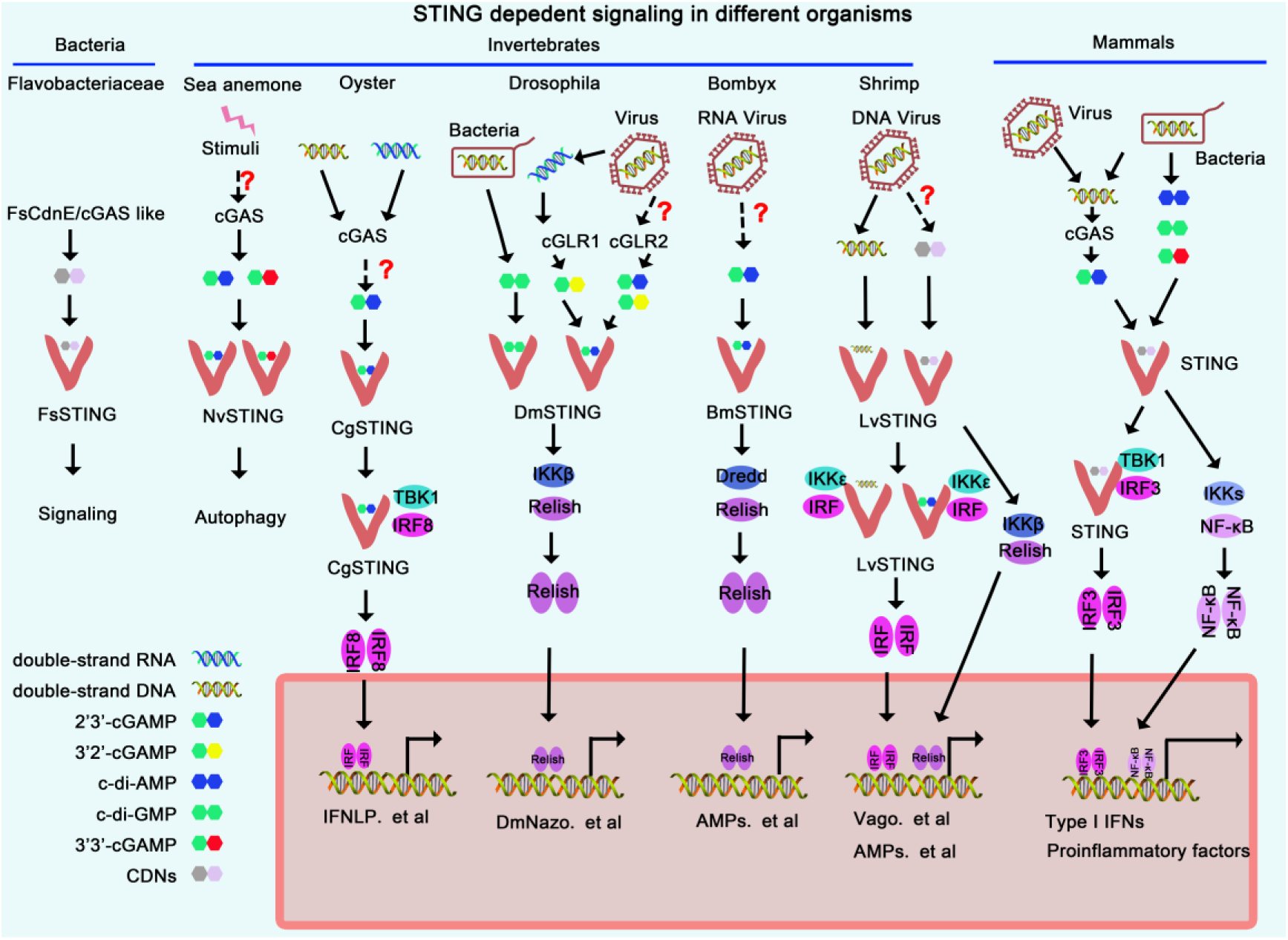
Diagram of STING dependent signaling from different species. The diagram of shrimp STING dependent immune signaling was mainly based on this study and previous reports [38, 55, 58, 60, 61, 66, 82, 83]. A mechanism of STING-dependent immune signaling in shrimps was proposed. In the mechanism, shrimp STING can directly sense DNA derived from pathogens, including DNA viruses; the activated STING then recruits IKKε and IRF and induces the activation of IRF transferred to the nucleus, where the antiviral factor Vago is expressed. Shrimp STING can be activated by bacterial infection and then induces NF-κB-mediated AMP production [38, 83]. The diagram of STING-dependent immune signaling in bacteria, sea anemone, oyster, insects, bombyx, and humans were based on previous reports [55, 58, 60, 61, 66, 82]. Question marks show that the mechanisms of activation of these steps are not well understood.

Owing to the loss of a zinc ribbon domain in cGAS, invertebrate cGAS cannot function as a DNA sensor. Therefore, invertebrate STING-dependent signal pathways may perform different activation mechanisms (Figure 9). In *N. vectensis*, NvcGAS produces cGAMP with a canonical 2′3′ linkage, and NvSTING activates antiviral autophagy through a mechanism that is independent of IFN induction in response to stimulation by the 2′3′-cGAMP produced by NvcGAS [57]. Interestingly, the NvcGAS activity of CDN synthesis can be activated by some unknown ligands, instead of DNA. This result indicated the presence of an ancient cGAS–CDN–STING pathway in *N. vectensis* that is not related to cytosolic DNA sensing [58]. An interesting divergence of STING dependent signaling was observed in Lophotrochozoa. This phylum, including *C. gigas* and *Capitella teleta*, contains several STING homologs with unusual structural architecture with TIR domains. TIR-domain-containing proteins are usually involved in Toll pathways, suggesting some associations between STING and Toll pathways in Lophotrochozoa [33]. In insects, STING-dependent pathways mainly emit signals through NF-κB activation [59]. For example, cGAMP is produced in silkworm *Bombyx mori* cells infected by a DNA virus named nucleopolyheedrovirus. *B. mori* STING (BmSTING) interacts with cGAMP and promotes the cleavage and nuclear translocation of NF-κB transcription factor Relish for efficient antiviral signaling [60]. *Drosophila* STING-dependent signaling performs different strategies against bacterial and viral infection. DmSTING senses c-di-GMP and induces antimicrobial peptides production to control *Listeria monocytogenes* infection through transcription factor Relish [61]. Liu *et al.* showed that DmSTING served as the target gene of NF-κB Relish and DmSTING induced antiviral autophagy to restrict Zika virus infection in the fly brain [62]. *Drosophila* cGAS-like protein DmcGLR1 can sense viral dsRNA and produce 2′3′-cGAMP, while DmGLR2 produces 2′3′-cGAMP and 3′2′-cGAMP in response to viral infection [63]. DmSTING can bind to 2′3′-cGAMP and 3′2′-cGAMP and then triggers the STING–IKKβ–NF-κB (Relish) signaling axis that contributes to opposing virus infection in flies [64–66]. All these studies about cGAS–STING signaling in invertebrates mainly focused on the relationships between CDN binding and Relish-mediated host defense. The mechanism underlying invertebrate cytosolic DNA-sensing induced STING–dependent IRF activation has not been thoroughly explored. Unlike CDNs mainly produced by bacteria, DNA exists in all pathogens, except in RNA viruses. Therefore, DNA is the most universal and conservative PAMP. The shrimp STING capable of DNA sensing might be a creative step in immune system evolution and connects direct DNA sensing from pathogens with STING–IRF triggered antiviral immune response.

In summary, we first demonstrated that an invertebrate STING can directly sense dsDNA to activate the IKKε–IRF–Vago antiviral signaling axis in a marine invertebrate. This feature indicates the ancient antiviral role of STING in innate immune responses, providing some novel insights into the functional origin of DNA sensing pathway in evolution.

## MATERIALS AND METHODS

The materials used are listed in Supplementary Table S1

### Animals and pathogens

Healthy shrimp (5 ± 0.5 g) were obtained from Guangdong Hisenor Group Co., Ltd Guangzhou, P. R. China. The shrimp were cultured in aerated seawater (30‰ salinity, 25 °C) of a recirculating aquaculture system with a water filtration plant, automatic temperature control equipment, and an ultraviolet sterilizer. Water was totally changed every 12 h. The commercial food (HAID Group) was supplied to the shrimp three times each day, and the leftover feed and excretions were removed in time. The healthy shrimp acclimated for three days before the injection experiments. Before all experiment treatments, shrimp (5% of total) were detected and confirmed to be free of common pathogens including white spot syndrome virus (WSSV), yellow head virus, taura syndrome virus, decapod iridescent virus 1, infectious hypodermal and hematopoietic necrosis virus and *Vibrio parahaemolyticus* by PCR or RT-PCR methods according to standard operation procedures by Panichareon *et al.* [67] and Qiu *et al.* [68]. Because WSSV and DIV1 were the main DNA viruses threatening shrimp culture, they thus were used here as potential activators for the shrimp STING-IRF pathway. WSSV particles were extracted from the muscle tissue of freshly WSSV-infected shrimp and stored at −80 °C according to a published method [69]. DIV1 particles were extracted from the cephalothorax tissue of DIV1-infected shrimp and stored at −80 °C according to a published method [70]. Before injection, a WSSV or DIV1 inoculum was prepared to approximately 1 × 10^5^ copies in 50 μL PBS. In the pathogenic challenge experiments, each shrimp received intraperitoneal injection of 50 µL of WSSV or DIV1 inoculum at the second abdominal segment with a 1-mL syringe.

### Primary hemocyte culture

The method for establishing primary cultured hemocytes was described in our previous study [71]. In brief, hemocytes from healthy shrimp *L. vannamei* were collected, suspended in serum–free Leibovitz-15 medium (Hyclone), seeded in 25 cm^2^ bottles, and maintained at 27 °C for 30 min. Then, the previous culture medium was replaced with fresh L15 medium containing 15% fetal bovine serum, 200 IU/mL penicillin and 200 μg/mL streptomycin.

### dsDNA synthesis and stimuli

Biotin-labeled, Cy5-labeled or unlabeled dsDNA named dsDNA1 or dsDNA2 were synthesized by Invitrogen (Thermo). The sequences are listed in Supplementary Table S2.

2′3′-cGAMP (0.2 μM) and dsDNA (2 μg/mL) was transfected into cells using FuGENE HD Transfection Reagent (Promega), whereas the transfection reagent without dsDNA or 2′3′-cGAMP was added as the control. WSSV or DIV1 was added to hemocytes cultured in medium with the concentration of 10^6^ copies/mL. After 6 h post challenge, the hemocytes were collected for quantitative reverse transcription PCR (qRT-PCR), Western blotting and immunofluorescence.

### Bioinformatics

To obtain the cDNA sequence of *STING* from shrimp, the amino acid (aa) sequences of the STING homologs from *Drosophila* (Genbank No. NM_001299327.1) and *Human* (Genbank No. NM_198282) were collected and used as query sequences for the *in silico* search of *L. vannamei* transcriptome data [72] by using local TBLASTN alignment tool with E-value cutoff of 1e^−5^. Specific primers (Supplementary Table S2) were designed based on the corresponding cDNA sequences, and were used for amplification of the *LvSTING* genome. The final genomic sequence was gained by overlapping with their adjacent fragments.

Genome and transcript sequences of STING homologs from other species were retrieved from the National Center for Biotechnology Information (NCBI, http://www.ncbi.nlm.nih.gov/). Sequence alignments were analyzed using Clustal X v2.0 [73]. Phylogenetic trees were constructed according to the deduced amino acid sequences with MEGA 5.0 software. Amino acid substitution type and Poisson model and bootstrapping procedure with a minimum of 1000 bootstraps were used [74]. The simple modular architecture research tool (SMART, http://smart.embl-heidelberg.de) was used in analyzing the deduced amino acid sequences of STING homologs [75]. The HMMTOP (http://www.enzim.hu/hmmtop/) was used in predicting LvSTING’s transmembrane domains [76].

3D structure models for the ligand-binding domains (LBDs) domains of LvSTING, GgSTING and HsSTING were generated with SWISS-MODEL (https://swissmodel.expasy.org/) [77]. The PDB for structural template identified for the LBD domains of LvSTING, GgSTING and HsSTING was 6nt7.1 (sequence identity of 23.29%, 97.96% and 79.03%, respectively). The 6nt7.1 describes the Cryo-EM structure of full-length chicken STING in the cGAMP-bound dimeric state [31].

### Plasmids construction

For *Drosophila* cell expressing system, the coding sequences of *LvSTING* (KY490589), *LvcGAS* (LOC113813773), *HscGAS* (KC294566), *LvIRF* (KM277954) and *LvIKKε* (AEK86520) were cloned into *Drosophila* S2 cell expressing plasmids for different tag-labeled proteins. The full-length or different segments of *LvSTING*, *LvIRF* and *LvIKKε* were amplified through PCR with corresponding primers listed in Supplementary Table S2, and cloned into the pAc5.1A-V5 (Invitrogen), pAc5.1A-HA [78], or pAc5.1A-GFP [79] vectors for expressing V5, HA, or GFP tagged proteins. The expression plasmids encoding the sequences of the deletion mutant of LvSTING were constructed through overlap extension PCR and cloned into pAc5.1A-HA. The primers used are listed in Supplementary Table S2.

### Prokaryotic protein expression and purification

The recombined cytoplasmic domain (CTD, 114-316) of LvSTING, and the CTD (137-376) of HsSTING were separately cloned into pMAL-c2x vectors (New England Biolabs), which were transformed into Rosseta cells for expressing MBP-tagged proteins. The MBP tagged proteins were purified by using Amylose Resin (New England Biolabs) and Ultrafiltration tubes (Millipore) according to the user’s manual.

### Pull-down assay

In the detection of the potential ability of different STING homologs to bind to 2′3′-cGAMP, 3′3′-cGAMP, c-di-AMP, c-di-GMP and DNA, several affinity gels, including c[8-AET-G(2’,5’)pA(3’,5’)p]-agarose (Biolog), c-(Ap-8-AET-Gp)-agarose (Biolog), 8-AET-c-diAMP-agarose (Biolog), and 8-AET-c-diGMP-agarose (Biolog) were used in the Pull-down assays. After equilibration with about 10 column volumes of starting buffer, the affinity column was loaded with the purified protein diluent or cell lysis. Non-specific binding proteins were removed as follows: c[8-AET-G(2’,5’)pA(3’,5’)p]-agarose was washed with washing buffer (10 mM cAMP and 10 mM ATP), c-(Ap-8-AET-Gp)-agarose was washed with washing buffer (10 mM cAMP and 10 mM ATP), 8-AET-c-diAMP-agarose was washed with washing buffer (10 mM GTP, 1 mM ATP, 0.25 mM cAMP and 0.25 mM cGMP), and 8-AET-c-diGMP-Agarose was washed with washing buffer (10 mM ATP, 1 mM GTP, 0.25 mM cGMP and 0.25 mM cAMP). Elution was performed with free 2′3′-cGAMP (100 μM; InvivoGen), 3′3′-cGAMP (100 μM; InvivoGen), c-di-AMP (100 µM; InvivoGen), and c-di-GMP (100 µM; InvivoGen). To determine whether STING homologs can bind to CDNs without non-specific electrostatic interactions, we incubated the monoclonal Dynabeads MyOne Streptavidin T1 (Thermo) was incubated with purified protein diluent and biotin (APExBIO) or biotin-poly (I:C) (InvivoGen) for 3 h at 4 °C, washed with PBS five times, and finally sampled with protein loading buffer. Pull-down assays were performed using SDS-PAGE, and Coomassie brilliant blue (CBB) staining was conducted. Pull-down assays for endogenous or ectopically expressed STING binding to dsDNA were performed with monoclonal Dynabeads MyOne Streptavidin T1. The beads were incubated with purified protein diluent or cell lysis containing endogenous STING (primary cultured shrimp hemocytes) or ectopically expressed STING, HscGAS or LvcGAS (*Drosophila* S2 cells) which transfected with biotin-dsDNA1/2 or biotin-poly (I:C) for 3 h at 4 °C, and then washed with PBS five times, and finally sampled with protein loading buffer. All the samples were analyzed with SDS-PAGE for CBB staining and Western blotting with antibodies of anti-actin (Sigma), anti-LvSTING (Genecreate), and anti-HA (Sigma).

### Co-IP

The interactions of endogenous proteins in shrimp hemocytes (*in vivo*) or ectopically expressed proteins in cells were investigated using Co-IP assays. Endogenous LvSTING–LvIRF interaction induced by viral infection was detected in hemocytes harvested 0, 12, and 24 h after WSSV or DIV1 infection. The hemocytes were lysed in IP lysis buffer (Pierce) with Halt Protease Inhibitor Cocktail (Thermo), and then incubated with protein G agarose beads (CST) coated with anti-LvSTING antibody (Genecreate) or a normal rabbit IgG antibody (CST) for 3 h at 4 °C, and finally detected by Western blotting with anti-LvIRF antibody (Genecreate). Five percent of lysed cells were loaded as input controls.

In the investigation of the interaction among proteins, the expression plasmids of the full length or deletion mutations of LvSTING, LvIKKε, and LvIRF with different tags were constructed. In brief, 48 h after transfection, *Drosophila* S2 cells were harvested and washed with ice-cold PBS three times and then lysed in IP lysis buffer (Pierce) with Halt Protease Inhibitor Cocktail (Thermo). The supernatants (500 µL) were incubated with 30 µL of anti-GFP (MBL), anti-V5 (Sigma) or anti-HA affinity gel (Sigma) at 4 °C for 4 h. The agarose affinity gels were washed with PBS five times and subjected to SDS-PAGE assay and Western blotting using rabbit anti-GFP antibody (Sigma), rabbit anti-V5 antibody (Merck) or rabbit anti-HA antibody (Sigma). Five percent total cell lysis was examined as the input controls.

### Western blotting

The hemocytes of virus-challenged, nonchallenged, DNA stimulated, and unstimulated shrimps were corrected and pooled from five shrimps for each sample at different time points. The nuclear and cytoplasmic fractions of hemocytes were extracted according to the protocol of NE-PER Nuclear and Cytoplasmic Extraction Reagents (Thermo). The total proteins were extracted by IP lysis buffer (Pierce) with a protease and phosphatase inhibitor cocktail (Merck), and then centrifuged at 12,000 *g* for 10 minutes at 4 °C to remove cell debris. For SDS-PAGE, protein samples were added with 5 × loading buffer and boiled for 10 min, and then subjected to Western blotting. For native PAGE protein samples were analyzed using 5% acrylamide gel (without SDS). Briefly, the gels were pre-run with 25 mM Tris and 192 mM glycine, pH 8.4, with and without 1% deoxycholate (DOC) in the cathode and anode chamber, respectively, for 30 min, at 50 mA. Samples in the native sample buffer (62.5 mm Tris– Cl, pH 6.8, 15% glycerol and 1% DOC) were applied to the gel and electrophoresed for 60 min at 50 mA, and further detected by Western blotting using anti-LvSTING or anti-LvIRF antibodies (Genecreate).

### qRT-PCR

The mRNA levels of genes in the pathogenic challenge experiments or *in vivo* RNA interference (RNAi) experiments were assessed with qRT-PCR assays. The expression levels of *LvSTING* and *LvVago4* were detected using a LightCycler 480 System (Roche, Basel, Germany) in a final reaction volume of 10 μL, which comprised 1 μL of 1:10 cDNA diluted with ddH_2_O, 5 μL of GoTaq qPCR Master Mix (Promega), and 250 nM primer (Supplementary Table S2). The cycling program was as follows: one cycle of 95 °C for 2 min, followed by 40 cycles of 95 °C for 15 s, 62 °C for 1 minute, and 70 °C for 1 s. Cycling was ended at 95 °C with a 5 °C/s calefactive velocity to create the melting curve. The expression level of each gene was calculated using the Livak (2^-ΔΔCT^) method after normalization to *EF*-*1a* (GU136229). Primers are listed in Supplementary Table S2.

### RNA interference (RNAi)

T7 RiboMAX Express RNAi System kit (Promega) was used to generate dsRNA-LvSTING dsRNA-LvcGAS and dsRNA-GFP (for controls) with primers containing a 5’ T7 RNA polymerase binding site (Supplementary Table S2). dsRNA quality was checked after annealing by gel electrophoresis. Each shrimp received an intraperitoneal injection of dsRNA at the second abdominal segment (2 μg/ g shrimp in 50 µl of PBS) or equivalent volume of PBS. Hemocytes were collected from shrimp 48 h after dsRNA injection, and total RNA was extracted and assessed by qRT-PCR with the corresponding primers for the evaluation of RNA interference (RNAi) efficacy. For survival experiment, 48 h after dsRNA-LvSTING or dsRNA-GFP injection, the shrimps were injected again with 1 × 10^5^ copies of WSSV or DIV1 particles or mock-challenged with PBS as a control. The survival rate of each group was recorded every 4 h 5–7 days after WSSV or DIV1 infection. The log-rank test method (GraphPad Prism software, GraphPad, USA) was used in analyzing differences among groups.

### Detection of viral loads by absolute quantitative PCR

Viral titers in shrimps were evaluated through absolute quantitative PCR. Briefly, we collected hemocytes from shrimp 48 h after WSSV or DIV1 infection. Hemocyte DNA was extracted with Marine Animal Tissue Genomic DNA Extraction Kit (TianGen). The quantity of WSSV genome copies was measured by absolute q-PCR using the primers of WSSV-F/R and a TaqMan fluorogenic probe named TaqMan probe-WSSV via LightCycler TaqMan Master kit (Roche) as described previously (Supplementary Table S2) [80]. Similarly, the primers of DIV1-F/R and TaqMan probe-DIV1 were used in obtaining the quantity of DIV1 genome copies (Supplementary Table S2) [81]. The WSSV or DIV1 genome copies were calculated and normalized to 1 µg of shrimp tissue DNA.

### Microscale Thermophoresis (MST)

MBP and MBP-LvSTING-CTD purified proteins were used to investigate their affinities to dsDNA through Microscale Thermophoresis (MST). Each purified protein, including MBP-LvSTING-CTD, MBP-HsSTING-CTD, MBP-DmSTING-CTD or MBP (as a control), was added in 1:1 dilution at 100 µM. Cyanine5 (Cy5)-labeled dsDNA1 was adjusted to the concentration of 20 nM. Samples were prepared in a buffer containing 20 mM Tris-HCl (pH 7.4), 1.5 mM MgCl_2_, 0.5 mM EDTA, 200 mM KCl, 10% glycerol and 0.05% NP-40 (v/v). Standard capillaries were filled with the samples for measurements with NanoTemper Monolith NT.115 instrument (NanoTemper, Germany). The measurement was performed in standard capillaries (NanoTemper, MO-Z022, Germany) at 20% LED and 50% MST power with laser-On time 30 sec and laser-off time 5 s.

### Dual luciferase assay

*Drosophila* S2 cells were cultured in a 24-well plate for dual luciferase assay. The cells of each well were transfected with 0.2 µg of Firefly luciferase reporter-gene plasmids (promoter region of LvVago4) [28], 0.04 µg of pRL-TK Renilla luciferase plasmids (internal control, Promega), and 0.01, 0.05 or 0.1 µg of protein expression plasmids or 0.5 µg of pAc5.1-V5 plasmids (as a control). Then, 48 h after -transfection, the cells were treated by 2′3′-cGAMP and dsDNA for 6 h. The cells were lysed, and 60% of the lysis was used in measuring Firefly of Renilla luciferase activity with Dual-Glo Luciferase Assay System (Promega). The remaining 40% of the cell lysis was analyzed through Western blotting, and the expression levels of proteins were detected. All the experiments were repeated three times.

### Immunofluorescence and confocal laser scanning microscopy

Hemocytes with different treatments, including viral infection, DNA transfection, and knockdown of specific genes, were spread onto coverslips in a 24-well plate. After 30 min, PBS was removed, and cells were fixed in 4% paraformaldehyde (diluted in PBS) at 25 °C for 15 min. The cells were then permeabilized with methanol at −20 °C for 10 min. After washing the slides three times, the hemocytes were blocked with 3% bovine serum albumin (diluted in PBS) for 1 h at 25 °C and then incubated with a mixture of primary antibodies (1:100, diluted in blocking reagent) overnight (about 8 h) at 4 °C. The primary antibodies used in immunofluorescence (IF) were rabbit anti-LvIRF antibody (Genecreate) and mouse anti-β-actin antibody (Sigma). The slides were washed with PBS six times and then incubated with diluted second antibodies of anti-rabbit IgG (H+L; 1:1000), F (ab′)2 fragment (Alexa Fluor 488 Conjugate; CST), and anti-mouse IgG (H+L), F (ab′)2 fragment (Alexa Fluor 594 Conjugate; CST) for 1 h at 25 °C. The cell nuclei were stained with Hoechst 33258 solution (Beyotime) for 10 min. Finally, after washing six times with PBS, the slides were observed with a confocal microscope (Leica, TCS-SP5, Germany).

### ChIP assay

Hemocytes from shrimp that were infected with WSSV after 24 h were used for ChIP assays. The ChIP assays were performed using a simple ChIP enzymatic chromatin IP kit (CST) according to the manufacturer’s instructions. In brief, hemocytes were cross-linked first with formaldehyde. Then, chromatin was digested with micrococcal nuclease into 150–900 bp DNA or protein fragments. Antibodies specific to rabbit LvSTING antibody or normal rabbit IgG antibody (provided by the kit as controls) were added, and complex co-precipitates were performed for 4 h at 4 °C with rotation and was captured by protein G magnetic beads for 2 h at 4 °C with rotation. The pelleted protein G magnetic beads were washed with low salt buffer three times. Then, high salt buffer was added to the beads and incubated at 4 °C for 5 min with rotation. The chromatin DNA was eluted from the antibody or protein G magnetic bead complexes, and the cross-links were reversed. DNA was purified using a spin column. The resulting purified DNA was subjected to semiquantitative PCR with 25 cycles of amplification. Primers were designed to amplify the sequences of Hr1, Hr5, the promoter regions of *wsv069*, *wsv078*, *wsv079*, *wsv249*, *wsv403*, and the coding sequences of *wsv184*, *VP15*, and *VP28* (Supplementary Table S2). The purified DNA from WSSV-infected shrimp was subjected to semi-quantitative PCR as positive controls. The products from semi-quantitative PCR without templates were considered negative controls. The PCR products were analyzed using agarose gel electrophoresis.

### Statistical analysis

All the data were presented as mean ± SD. Student’s *t*-test was used in calculating the comparisons among the groups of numerical data. For survival rates, data were subjected to statistical analysis using GraphPad Prism software to generate the Kaplan ± Meier plot (log-rank χ^2^ test). The following *p*-values were considered statistically significant: **\****p* < 0.05, **\*\****p* < 0.01, and **\*\*\****p* < 0.001.

## DATA AVAILABILITY

All reagents and experimental data are available within Supplemental transparent methods or from corresponding author upon reasonable request.

## ACKNOWLEDGEMENTS

This research is supported by National Natural Science Foundation of China (32173000/32022085/31930113), Independent Research and Development Projects of Maoming Laboratory (2021ZZ007/2021TDQD004) and Southern Marine Science and Engineering Guangdong Laboratory (Zhuhai) (SML2021SP301), Science and Technology Planning Project of Guangzhou City (202102020354), and the Fundamental Research Funds for the Central Universities, Sun Yat-sen University (22lglj05). The funders have no role in study design, data collection and analysis, decision to publish, or preparation of the manuscript. The authors are grateful to Guangdong Hisenor Group Co., Ltd Guangzhou for providing specific pathogen free shrimp.

## AUTHOR CONTRIBUTIONS

Chaozheng Li and Jianguo He conceived and designed the experiments. Haoyang Li, Qinyao Li, Sheng Wang and Shaoping Weng performed the experiments and analyzed data. Haoyang Li, Jianguo He and Chaozheng Li wrote the draft manuscript. Chaozheng Li and Jianguo He acquired finding. All authors discussed the results and approved the final version.

---

**Figure 1–figure supplement 1.**
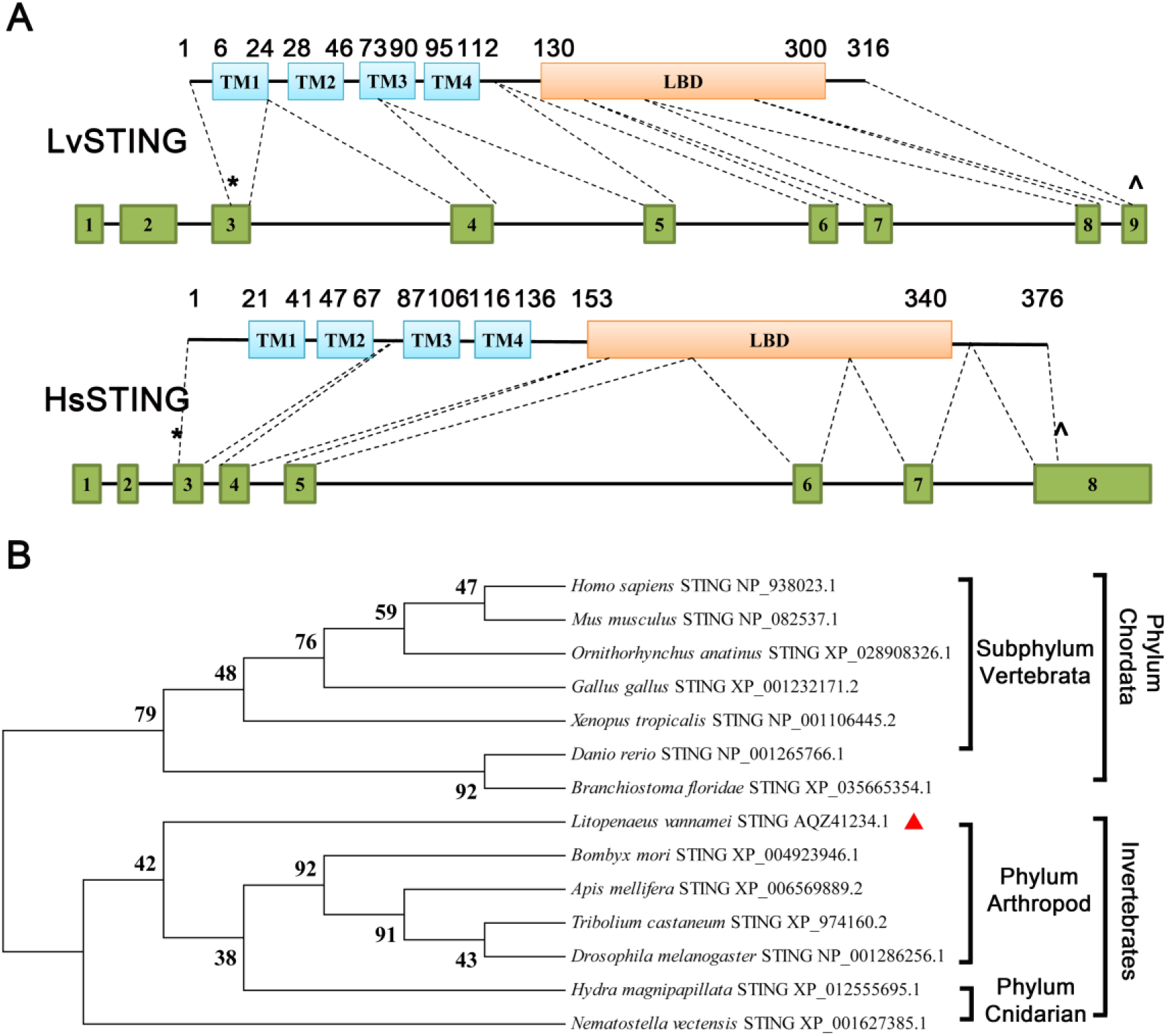
Comparison among STING homologs. **(A)** Schematic representation of the genomic structures of LvSTING and HsSTING. Exons (numbered boxes) are shown on the genomic sequence (horizontal line), and start and stop codons are indicated by asterisks (**\***) and carets (^), respectively. Numbers above the boxes indicated the amino acid start points of each putative domain. GenBank number of LvSTING genomic sequence was OP908242. **(B)** Phylogenetic tree of STING homologs. The tree was constructed with the neighbor-joining (NJ) method based on 14 STING full-length protein sequences aligned with MEGA 5.0. LvSTING was indicated in red diamond.

**Figure 3–figure supplement 1.**
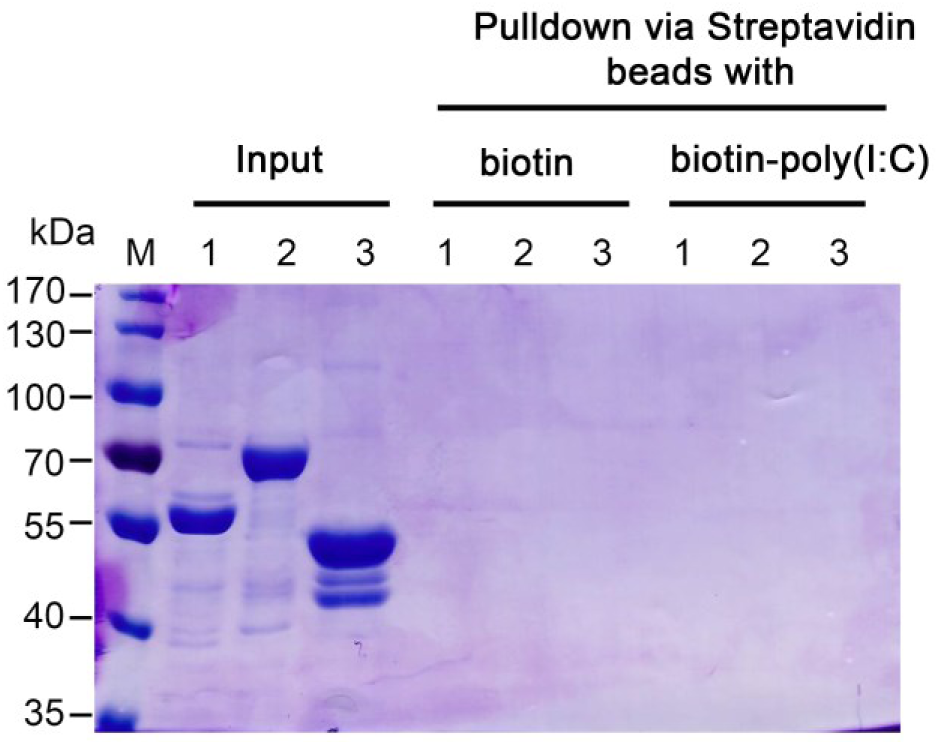
LvSTING could not interact with dsRNA. Pull-down assay to determine interaction between STING and biotin or biotin-poly (I:C). MBP-HsSTING-CTD, MBP-LvSTING-CTD and MBP were pulled down via streptavidin beads with biotin or biotin-poly (I:C). All experiments were replicated three times with similar results.

**Figure 4–figure supplement 1.**
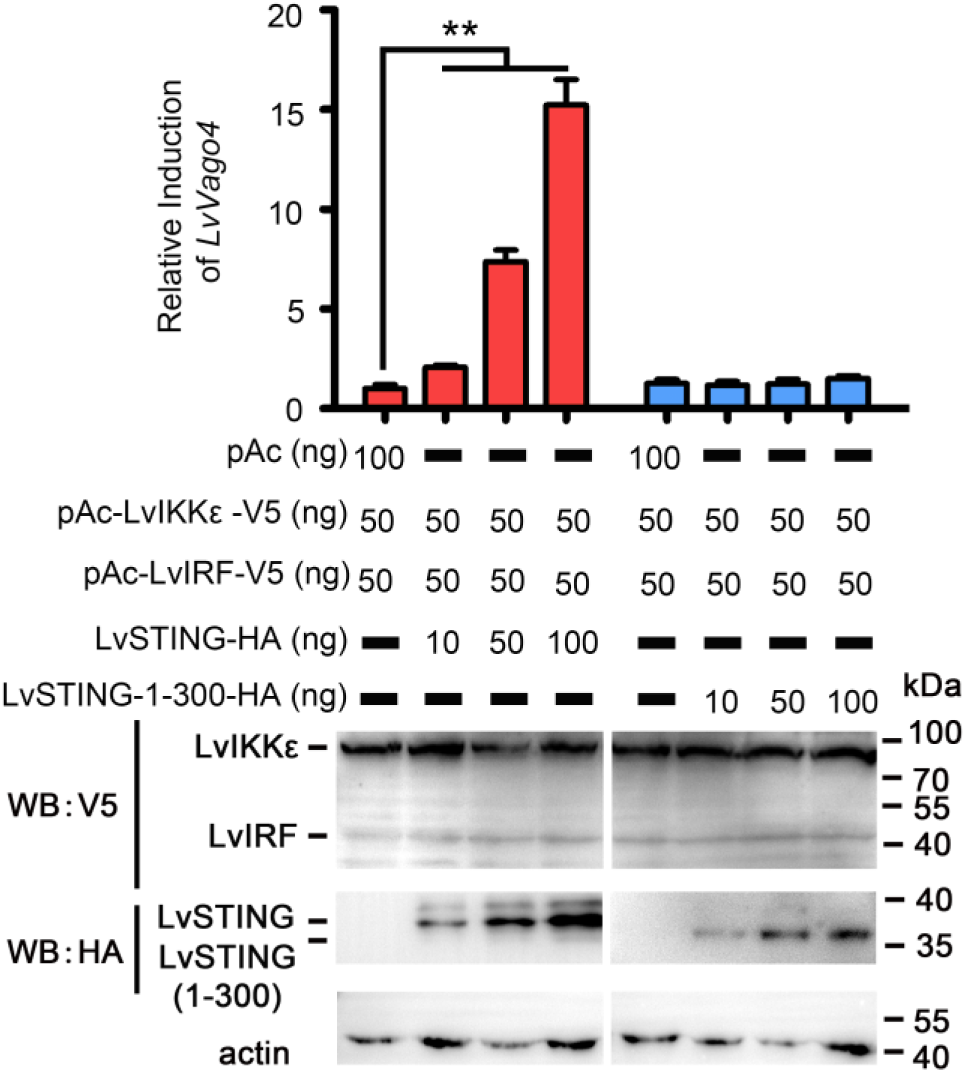
LvSTING-dependent LvVago4 activation was required for sCTT domain. Relative induction of the promoter activities of *LvVago4* mediated by ectopically expressed LvIRF, LvIKKε and LvSTING or LvSTING-1-300 in *Drosophila* S2 cells in dual luciferase assays. Ectopically expressed proteins were detected by Western blotting. All experiments were replicated three times with similar results.

**Figure 6–figure supplement 1.**
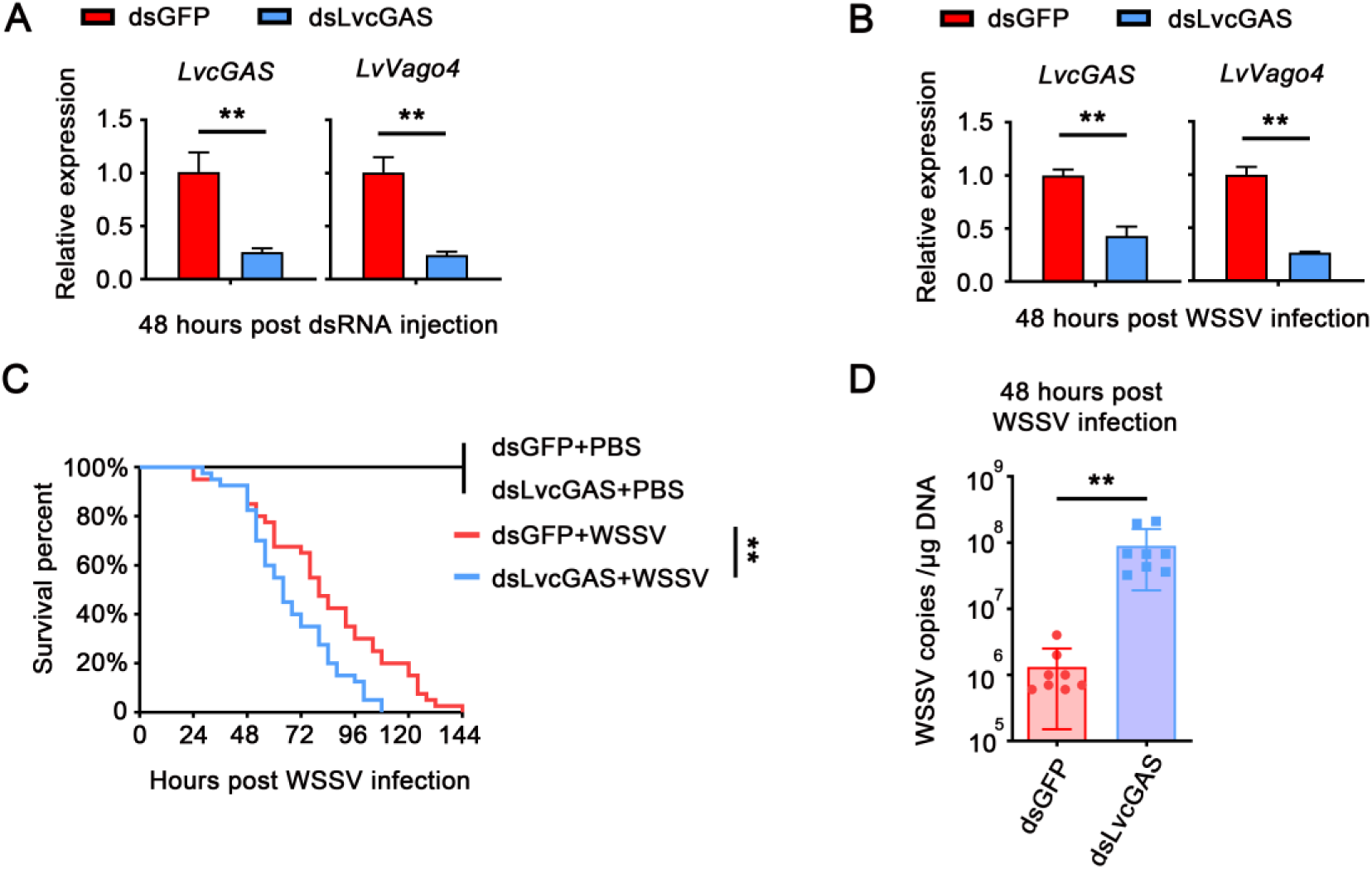
LvcGAS defended against WSSV infection. **(A)** Relative expression of LvcGAS or LvVago4 was detected 48 h after dsRNA-LvcGAS injection. **(B)** Relative expression of LvcGAS or LvVago4 was detected 48 h after WSSV infection. **(C)** Survival rates of WSSV infected shrimp injected with dsRNA-LvcGAS or dsRNA-GFP. Shrimp survival rate was monitored every 4 h after viral infection. Differences between groups were analyzed with log-rank test performed using GraphPad Prism 5.0 (**\*\****p* < 0.01, **\*\*\****p* < 0.001). **(D)** Virus titers in the hemocytes of shrimps treated with dsRNA-LvcGAS or dsRNA-GFP (as control) 48 h after WSSV infection. Student’s t-test was applied (**\*\****p* < 0.01). All experiments were replicated three times with similar results.

**Figure 8–figure supplement 1.**
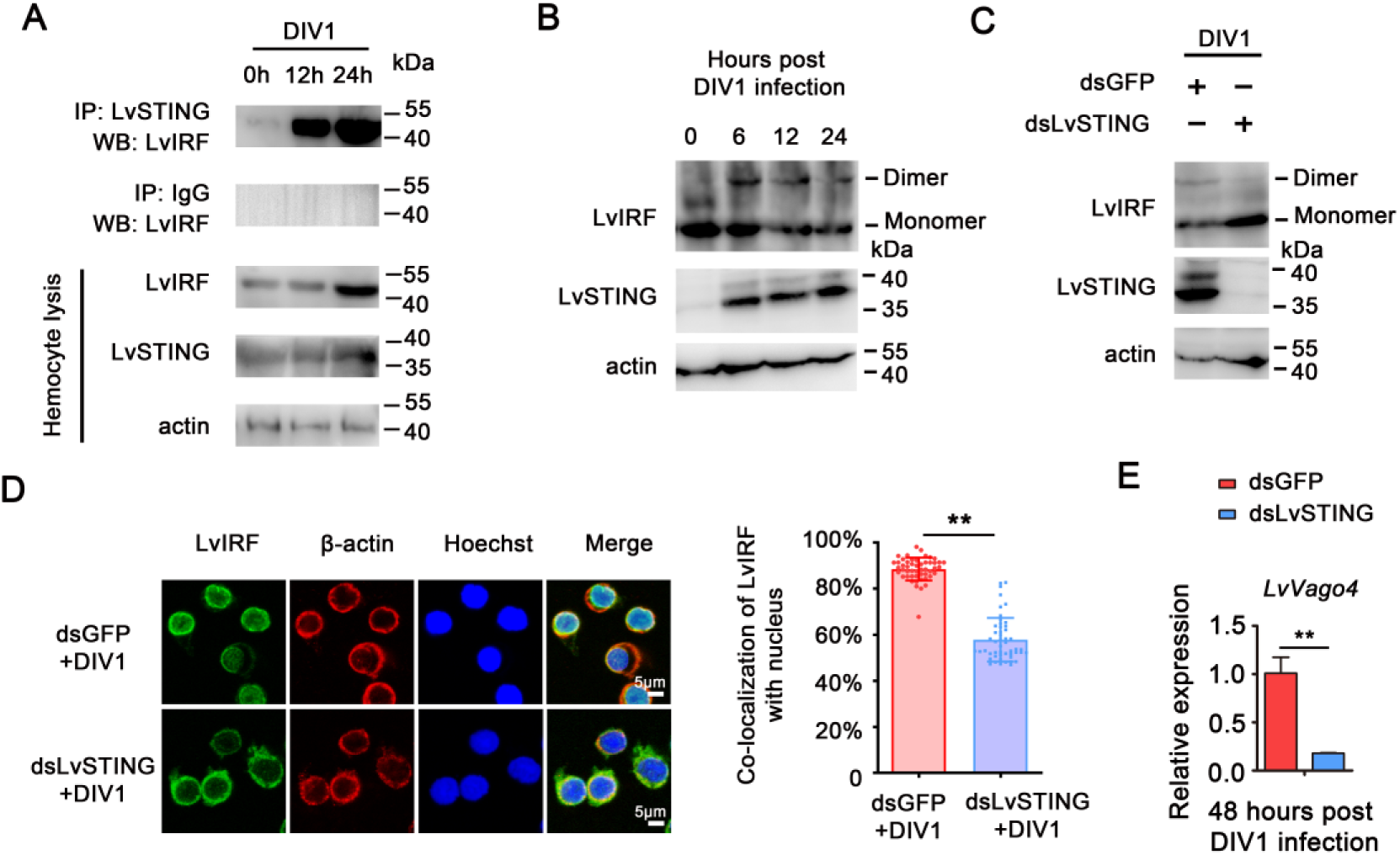
DIV1 infection activated the shrimp STING–IRF– Vago4 axis *in vivo*. **(A)** Increased interaction of LvSTING-LvIRF 0, 12, and 24 h post DIV1 infection *in vivo*. **(B)** Increased IRF dimerization 0, 12, and 24 h post DIV1 infection *in vivo*. **(C)** Knockdown of LvSTING attenuated the dimerization of LvIRF in shrimp hemocyte during DIV1 infection. **(D)** LvIRF nuclear translocation in response to DIV1 infection was inhibited by the knockdown of LvSTING. Co-localization of LvIRF and nucleus in hemocytes (*n* =50 cells) calculated by ImageJ and analyzed statistically by student’s *t*-test (**\****p* < 0.05). **(E)** Silencing of LvSTING attenuated the transcriptional levels of *LvVago4* in hemocytes during DIV1 infection. Data were analyzed statistically by student’s *t*-test (**\*\****p* < 0.01). All experiments were replicated three times with similar results.

